# Benchmarking computational methods for B-cell receptor reconstruction from single-cell RNA-seq data

**DOI:** 10.1101/2022.03.24.485600

**Authors:** Tommaso Andreani, Linda M. Slot, Samuel Gabillard, Carsten Struebing, Claus Reimertz, Yaligara Veeranagouda, Aleida M. Bakker, Reza Olfati-Saber, Rene’ E. M. Toes, Hans U. Scherer, Franck Augé, Deimantė Šimaitė

**Affiliations:** AI & Deep Analytics – Omics Data Science, Sanofi, Frankfurt am Main, 65926, Germany; Immunology & Inflammation Research, Sanofi GmbH, Frankfurt am Main, 65926, Germany; AI & Deep Analytics, Sanofi, Cambridge, MA, 02142, United States; AI & Deep Analytics – Omics Data Science, Sanofi, Paris, 91385, France; Molecular Biology & Genomics, Translational science unit, Sanofi, Chilly-Mazarin, 91385, France; Department of Rheumatology, Leiden University Medical Center, 2333 RC Leiden, The Netherlands; Life & Soft, Le Plessis-Robinson, Paris, 92260, France

## Abstract

Multiple methods have recently been developed to reconstruct full-length B-cell receptors (BCRs) from single-cell RNA-seq (scRNA-seq) data. This need emerged from the expansion of scRNA-seq techniques, the increasing interest in antibody-based drug development and the importance of BCR repertoire changes in cancer and autoimmune disease progression. However, a comprehensive assessment of performance-influencing factors like the sequencing depth, read length or the number of somatic hypermutations (SHMs) as well as guidance regarding the choice of methodology are still lacking. In this work, we evaluated the ability of six available methods to reconstruct full-length BCRs using one simulated and three experimental SMART-seq datasets. In addition, we validated that the BCRs assembled *in silico* recognize their intended targets when expressed as monoclonal antibodies. We observed that methods like BALDR, BASIC and BRACER showed the best overall performance across the tested datasets and conditions whereas only BASIC demonstrated acceptable results on very short read libraries. Furthermore, the *de novo* assembly-based methods BRACER and BALDR were the most accurate in reconstructing BCRs harboring different degrees of SHMs in the variable domain, while TRUST4, MiXCR and BASIC were the fastest. Finally, we propose guidelines to select the best method based on the given data characteristics.

## INTRODUCTION

The recent development of single-cell RNA sequencing (scRNA-seq) techniques has enabled the quantification of genes expressed in individual cells. This has contributed to the identification of different cell types characterizing tissues and to the discovery of previously unknown cell populations, in which the underlying gene expression programs were found to be critical for embryonic development, autoimmune disease pathogenesis and an in-depth understanding of the tumor microenvironment (1–4). Similarly, following the increasing evidence of the importance of B lymphocytes in health and disease, single-cell technologies have been applied to quantify the expression levels of genes, coding for heavy (HC) and light chains (LC) of a B-cell receptor (BCR) and thus the BCRs/antibodies these cells produce.

Human BCRs consist of a pair of independent HC and LC that are interconnected by disulfide bonds, each of which contains both constant (C) and variable (V) regions, genetically encoded in three different loci. The immunoglobulin heavy chain locus (IGH) on chromosome 14 contains gene segments for the immunoglobulin HC, whereas LC genes are encoded by two loci: the immunoglobulin kappa (κ) chain locus (IGK) on chromosome 2 and the immunoglobulin lambda (λ) chain locus (IGL) on chromosome 22. During the HC somatic recombination, one of the diversity (D) gene segments is joined to one of the joining (J) gene segments in an event called the D-J recombination (5). Afterwards, the D-J segment binds one of the variable (V) segments and all constant regions are retained at the end of the mRNA to produce a functional HC. Since LC do not have a D segment, only the V-J recombination occurs. During these processes, random nucleotides are added into the V(-D-)J joining regions, resulting in a higher number of HC and LC than it would be possible by simply joining gene segments available at each HC and LC genes’ locus. This process, in fact, can generate an immunoglobulin repertoire of more than 5×10¹³ different antigen specificities (5). The introduction of these new nucleotides is particularly challenging for BCR assembly algorithms because of the randomness of such introductions and the absence of them in the reference genome and in immunoglobulin annotation databases.

The characterization of BCR repertoires from scRNA-seq data has been instrumental in the investigation of groups of B-cells sharing a common ancestor, called clonotypes. An overrepresentation of pathogenic clonotypes was noticed in different diseases such as breast cancer (6), multiple sclerosis (MS) (7) and acute myeloid leukemia (AML) (8). In these studies, the presence of repertoires with expanded clonotypes defined different tumor microenvironments in breast cancer, were associated with increased inflammation in MS and could be used to stratify AML patients. In addition, BCRs are known to acquire somatic hypermutations (SHMs) in their variable domain (defined as four framework regions (FWRs) and three complementary-determining regions (CDRs)) after activation of the B-cell by an antigen. In the process of affinity maturation, the affinity for a given antigen can be enhanced by introducing somatic mutations predominantly in the CDR regions. For this reason, BCRs of memory B-cells display somatic hypermutation that are not germline-encoded when compared to naïve B-cells. As memory B-cells represent the cell population that has been triggered by a given antigen and hence are mostly connected to processes studied in e.g., autoimmunity or infectious diseases, these cells often gain most interest in studies investigating BCR composition. For example, an increasing number of SHMs in anti-citrullinated protein antibodies (ACPAs) during rheumatoid arthritis (RA) development (9) has been described together with a high frequency of defined sequences called N-glycosylation sites that can potentially be used as a predictor of RA progression (10). Furthermore, BCRs of patients with diffuse large B-cell lymphoma (DLBCL) harbor variable levels of SHMs in the variable (V) regions of IGH and IGL/IGK genes. Here, high levels of clonal IGHV SHMs were associated with a prolonged overall survival of patients whereas an increased CDR3 length of HC and the presence of IGHV ongoing SHM were associated with poor prognosis (11). Overall, these studies show the importance of patients’ BCRs in the pathogenesis of multiple diseases and the potential usage of their characterization as a proxy in personalized medicine.

Previous sequencing techniques targeting the immunoglobulin genes, such as Ig-seq (12), allowed the quantification of the entire set of genes belonging to the HC and LC in the total population of B-cells (also called BCR repertoire), giving a snapshot of the repertoire composition. However, the inability to obtain full-length BCRs consisting of the variable domain of a HC-LC pair set in an individual cell has been a limiting factor of such approaches. Nowadays, there are several different scRNA-seq technologies available. These can be plate-based, which are generally low throughput but can be used to sequence full-length transcripts, or droplet-based, allowing to sequence thousands of cells at the same time. Two plate-based approaches, namely, SMART-seq (13, 14) and a modification of it named SPEC-seq (15) have been instrumental in BCR sequencing and full-length BCR reconstruction due to their ability to obtain full-length transcripts of HC and LC genes allowing the reconstruction of the variable domain of a HC-LC pair set in a single cell. The droplet-based 10x Genomics Chromium Single Cell Immune Profiling Solution (16) also enables sequencing of the V-D-J genes of a B-cell to obtain paired HC and LC. However, this comes with the cost of losing the full-length BCR information for a considerable number of single cells (17).

Although it is not feasible to determine a pair of HC and LC constituting a BCR of a cell using bulk RNA-sequencing techniques, several methods have been proposed to delineate BCR composition from complex data sets. Methods, such as (18)(18)(19)(20–22) and Imonitor (23) are only suitable for the reconstruction of complementarity-determining region 3 (CDR3 of the variable domain of HC and LC. Therefore, BCRs, reconstructed using the above methods, miss CDR1, CDR2 regions and the four FWRs. Nonetheless, these regions considerably contribute to the antigen recognition and binding and aid in maintaining the overall structure of an antibody (24). For this reason, the analysis of BCR repertoires from scRNA-seq data required the development of algorithms capable of dealing with highly mutated sequences, dissimilar to the reference genome in order to reconstruct the full variable domain of HC-LC pair of a B-cell. Recently, several open-access methods have been proposed for preprocessing of raw SMART-seq and Chromium data to reconstruct BCRs (**Table 1**). The first developed method was MiXCR (25), which consists of a collection of algorithms based on a proprietary aligner that perform clustering to accomplish BCR reconstruction and annotation. BASIC, a semi *de novo* algorithm (26) was the second method that was made available with the advantage of being able to process libraries as short as 25 base pairs (bp). Lately, several algorithms based on a *de novo* assembly but using different approaches to map reads and assign V-D-J genes such as BRACER (27), BALDR (28), VDJpuzzle (29) and TRUST4 (30) have emerged, increasing the choice but also the difficulty in selecting the best tool for a given dataset. Importantly, MiXCR, VDJpuzzle and TRUST4 can be used for both BCR and T-cell receptor (TCR) assembly, making them suitable for more elaborate immune repertoire studies.

**Table 1.** Description of the computational methods for BCR reconstruction from scRNA-seq data evaluated in this benchmark.

| Method | Availability | Language | Type of Algorithm | Annotation | Species | Sequencing | Publication |
| --- | --- | --- | --- | --- | --- | --- | --- |
| MIXCR | https://github.com/milaboratory/mixcr | Java | Proprietary aligner using *k*-mers & assembler | IMGT | Human, Mouse | SMART-seq2 | Bolotin 2015 |
| BASIC | https://github.com/akds/BASIC | Python | Semi *de-novo* with anchors & *k*-mers | IMGT | Human, Mouse | SMART-seq2 | Canzar 2017 |
| BRACER | https://github.com/Teichlab/bracer | Python/R | *de-novo* | IMGT, Combinatorial Recombinome | Human, Mouse* | SMART-seq2 | Lindeman 2018 |
| BALDR | https://github.com/BosingerLab/BALDR | Perl | *de-novo* | IMGT, Combinatorial Recombinome | Human, Rhesus Macaque | SMART-seq2 | Upadhyay 2018 |
| VDJPuzzle | https://github.com/simone-rizzetto/VDJPuzzle | Roff/Shell | *de-novo* | IMGT | Human, Mouse | SMART-seq2 | Rizzetto 2018 |
| TRUST4 | https://github.com/liulab-dfci/TRUST4 | C++/Python | *de-novo* with *k*-mers | IMGT | Human, Mouse | SMART-seq2 + 10x | Song 2021 |

\* Possibility to obtain other species

Given the methodological differences in the algorithms (see **Supplementary Information**), they can report different results under certain experimental setups. Conditions, such as different sequencing technologies, read library properties and the number of SHMs expected within the variable domains of the BCRs can influence the output of the tools. Thus, it is essential to assess the performance of each algorithm and quantitatively understand how sensitive it is in reconstructing BCRs when compared to the “ground truth” (defined as the original variable domain sequence obtained by classical Sanger sequencing). In addition, given varying numbers of SHMs within the variable domains of BCRs, and the observation of B-cells with high SHMs counts in several diseases like follicular lymphoma (31), rheumatoid arthritis (9), diffuse large B-cell lymphoma (11) as well as in anti-HIV antibodies (32–34), it is crucial to assess the number of SHMs the algorithms can tolerate while still accurately reconstructing BCRs. Finally, the comparison of these methods using several datasets will reveal the stability of their performance across different library preparations.

To address these questions, we designed a comprehensive analysis framework (**Figure 1**). Firstly, we selected two publicly available BCR sequencing datasets of plasmablast origin with the available ground truth (26, 28) and generated an additional dataset with the ground truth, named “Leiden”, using sorted isotype-switched memory B-cells that either recognize tetanus toxoid (TT) or have an unknown antigen specificity. Secondly, using random sampling, we created multiple libraries with different levels of coverage and read length using the above-mentioned experimental datasets. Thirdly, we simulated a fully synthetic dataset (see **Material** and **Methods**) in which HC and LC harbor different amount of SHMs. Subsequently, we used these datasets to test six available BCR reconstruction methods. We evaluated their abilities to obtain productive HC and LC, defined by the absence of stop codons or out-of-frame V-J junctions (see **Material** and **Methods**). This was carried out using the sensitivity as a metric for the experimental data and accuracy as a metric for the synthetic data (see **Methods**). In addition, we experimentally validated the benchmarked methods by determining that antibodies, generated using the productive sequences, can recognize their intended antigens. Moreover, we measured the time needed to run these algorithms to provide information on time and scalability. Finally, we propose a final performance score for each method by aggregating the results of all the experiments and provide recommendations for the selection of the method that best suits a given dataset and scientific question.

**Figure 1.**
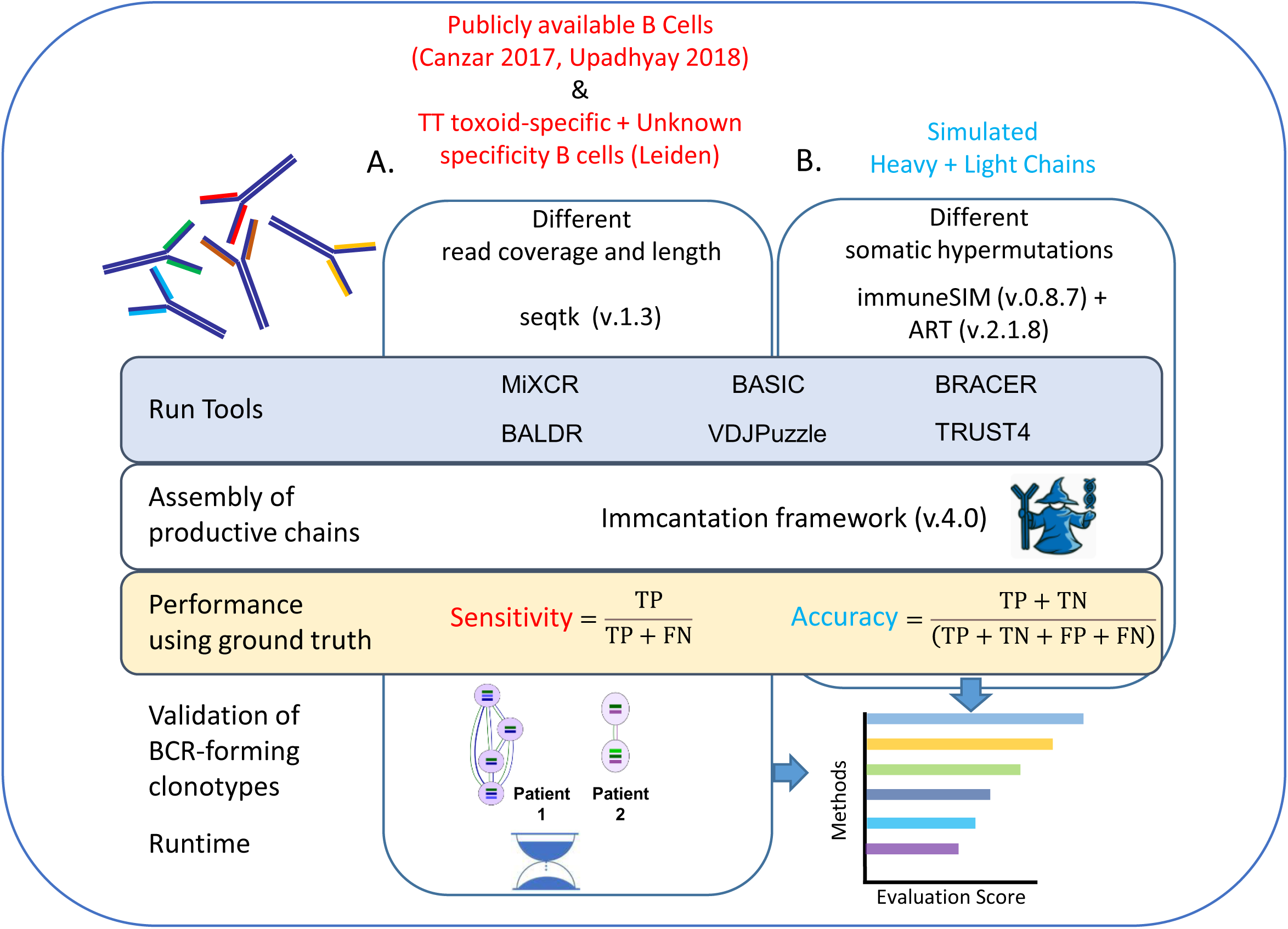
Benchmark framework: **(A)** Available datasets with ground truth (Canzar et al. 2017, Upadhyay et al. 2018) consisting of plasmablasts with unknown antigen specificity were obtained from the corresponding publications. In addition, a scRNAseq dataset including tetanus toxoid-specific B-cells and B-cells with unknown antigen specificity was generated in this work (Leiden). All the datasets were downsampled to achieve different read coverage and length. **(B)** An additional dataset was simulated to investigate the effects of different levels of somatic hypermutations in the variable domains of BCRs on the performance of each method. Sensitivity and accuracy were used as metrics to evaluate each method depending on the type of used data. Antibodies were produced using a subset of clonotype-forming patient-specific BCRs and their specificity was experimentally validated. Finally, the execution time was investigated, and a final score was calculated to give a final recommendation on the method choice.

## MATERIAL AND METHODS

### Experimental Datasets

#### Leiden dataset (SMART-seq2)

##### Cell sorting, cDNA synthesis, ARTISAN PCR and Sanger sequencing

TT-specific B-cells and B-cells with unknown specificity were isolated as described before (9). In short, peripheral blood mononuclear cells (PBMCs) were isolated by Ficoll-Paque gradient centrifugation and stained with Fixable Violet (405nm) Dead Cell Stain kit (Thermofisher), CD3 Pacific Blue (clone UCHT1, BD Pharmingen), CD14 Pacific Blue (clone M5E2, BD Pharmingen), CD19 APC-Cy7 (clone SJ25C1, BD Pharmingen), CD20 AlexaFluor 700 (clone 2H7, BD Pharmingen), CD27 PE-Cy7 (clone M-T271, BD Pharmingen), IgG BV510 (BD Horizon, clone G18-145), IgD FITC (clone IA6-2, BD Pharmingen) and APC- and PE-labeled tetanus toxoid (TT). CD19^+^CD20^+^CD27^+^IgG^+^IgD^-^ B-cells were considered TT-specific if they stained double positive for fluorescently labeled TT with two different fluorochromes; TT- APC and TT-PE. Cells negative for two labeled antigens were considered as ‘cells with unknown specificity’. Cells were single-cell sorted on a FACS ARIA sorter and mRNA was lysed directly in lysis mix; 0.2% Triton X-100 (Sigma) in ddH2O, RNase inhibitor (25 U, TaKaRa), Oligo-dT30VN (10 pmol, IDT) and dNTPs (10 nmol, ThermoFisher). cDNA synthesis and subsequent preamplification and purification was performed according to the SMART-seq2 protocol (14). Anchoring Reverse Transcription of Immunoglobulin Sequences and Amplification by Nested (ARTISAN) PCR was performed using purified cDNA.

The ground truth ARTISAN Sanger sequencing of each single cell was performed on an Applied Biosystems 96-capillary (ABI3730xl) sequencer. After counting the HC and the LC of the sequenced 72 single cells, 56 HC, 56 kappa light chains (LcK) and 27 lambda light chains (LcL) were defined as *assembled*. These were HC and LC that did not contain a stop codon in variable and constant regions as defined by IgBLAST and Change-O (35). Furthermore, 38 HC, 48 LcK, 8 LcL and 27 paired HC + LC (K or L) were classified as *productive*. IgBLAST and Change-O (35) define a chain *productive* if it is an *assembled* chain with in-frame V-J junctions. These 27 single cells with paired HC + LC were used as the ground truth to compute the sensitivity of each method for this dataset. (**Supplementary Figure S1A, Supplementary Table S3**).

#### Canzar dataset (SPEC-seq)

We retrieved the dataset included in the publication of BASIC (26) from GEO (https://www.ncbi.nlm.nih.gov/geo/query/acc.cgi?acc=GSE116500). Out of 295 single plasmablast cells published in this dataset, we selected 190 that were sequenced using a 50 base-pair paired-end mode. We further filtered for 113 samples that had at least 1.25 million reads. Besides the fastq files, we obtained fasta sequences generated using Sanger sequencing, which we used as the ground truth for this dataset. Afterwards, we ran the Immcantation framework v 4.0 (35) to assess the number of *assembled* and *productive* HC and LC (K or L) in each single cell and to determine germline genes in each sequence. As a result, we assembled 113 HC, 62 kappa light chains (LcK) and 57 lambda light chains (LcL), including 102 cells having complete HC+LC pairs. After the assembly, 68 HC, 48 LcK and 46 LcL, including 45 paired HC + LC (complete B-cell receptors), were labeled as *productive*. Hence, we used these 45 single cells with full-length paired and productive HC and LC to compute the sensitivity of each method for this dataset. (**Supplementary Figure S1B**)

#### Upadhyay dataset (SMART-seq2)

We retrieved the human AW1 plasmablast dataset included in the BALDR publication (28) from SRA (https://www.ncbi.nlm.nih.gov/sra/?term=SRP126429). This dataset, consisting of 51 single cells, was firstly investigated for the read quality using fastQC v0.11.9 (36). During this quality control step, we observed that libraries contained duplicated reads, overrepresented sequences and *k*-mers despite the quality of the reads being adequate at 3’ and 5’ ends (**Supplementary Data 1** and **Supplementary Figure S3)**. After downloading fastq files and fasta sequences of the ground truth that were generated using Sanger method, we ran the Immcantation framework v 4.0 (35) to assess the number of *assembled* and *productive* HC and LC in each single cell and to determine germline genes in each sequence. As a result, we reconstructed 34 HC, 19 kappa light chains (LcK) and 22 lambda light chains (LcL), including 23 paired HC + LC that were termed as *assembled*. Out of these, 34 HC, 19 LcK and 21 LcL were labeled *productive*, including 23 HC + LC pairs (B-cell receptors). Consequently, we used these 23 single cells with full-length paired and productive HC and LC to compute the sensitivity of each method for this dataset (**Supplementary Figure S1C**).

#### Simulation of experimental datasets with different read length and coverage

We used seqtk (37) version 1.3-r106 with the option ‘-s100’ to perform random sampling without reintroduction of additional reads to generate libraries with different levels (from 50,000 up to 1.25 million reads) of coverage. Afterwards, the sampled reads were trimmed using seqtk with the option ‘trimfq’ to obtain final libraries with read lengths ranging from 25 up to 50 nucleotides for the Canzar dataset (26), and from 25 up to 100 nucleotides for the Upadhyay dataset (28) and for the Leiden dataset. In total, we simulated 70 distinct libraries covering different read length, coverage, type of B-cells (memory and plasmablasts) and plate-based techniques (SMART-seq2 vs SPEC-seq).

#### Simulation of synthetic chains with different levels of somatic hypermutations

The process of generating SHMs has been described as stochastic in nature for many decades. However, it has recently been demonstrated that intrinsic biases are present *in vivo* mostly due to the activity of the activation-induced cytidine deaminase (AID) (38). Nevertheless, we wanted to test whether a mutation accruing at any position of the CDR in the variable domain could generalize an effect on the performance of the methods used in this benchmark. For this, we used immuneSIM (39) version 0.8.7 to simulate four libraries containing 100 HC, 100 LcL and 100 LcK that harbored 15, 30, 45 and 60 (SHMs). We used the option “shm.mode=data”, which focuses on mutation events in the CDRs (based on IMGT) occurring at any position during the process of SHM. Firstly, this tool computes the frequencies of the V-D-J germlines genes, insertions and deletions in the V and J junctions using repertoires present in different studies (40–43). Secondly, it recreates the sequences of HC and LC trying to maintain the statistical patterns and ratios of the number of germline genes. In case an introduction of SHMs is required in a simulation, the tool considers only chains without stop codons. In case a stop codon occurs, it resamples genes and other features until the requested number of simulated sequences without stop codons is reached. We noticed, though, that even if the resulting simulated sequences of HC or LC had no stop codons, they could still have out-of-frame V-J which eventually resulted in a non-productive chain. The output fasta file of each of the simulated HC, LcL and LcK was used as a reference to create synthetic reads and to obtain paired-end Illumina libraries containing 500,000 75 bp long reads, using ART (44) version 2.1.8 (parameters *-l* 75, *-f* 500,000). Finally, we annotated each of the simulated sequences using the Immcantation framework v 4.0 (35) to identify the germline genes and the number of *assembled* and *productive* HC and LC (**Supplementary Figure S2 A** to **D**, **Supplementary Table S4**) which we used as the ground truth to compute the accuracy of each tool.

#### Sensitivity as a performance metric for experimental data

In this study, ground truth was considered as a set of HC and LC sequences coming from different single cells and obtained by Sanger sequencing. In principle, every cell should have a pair of HC and LC in the ground truth. However, we realized that only ∼40% of single cells in each of three analyzed datasets had paired HC and LC that were also defined as *assembled* and *productive* using the criteria as in the chapters above (**Supplementary Figure S1 A to C**). Therefore, the ground truth information was available only for a subset of single cells (and thus productive BCRs) that could be used as a ground truth for all the single cells in each dataset. A metric such as specificity, could not be used to evaluate the performance of the tested algorithms for BCR reconstruction due to the nature of the data (every antigen-specific B-cell must have a HC and a LC) which prevents calculation of the false positive values. In other words, if Sanger sequencing misses one HC or one LC, assigning a false positive to a productive HC and LC assembled by a tool would be a wrong assumption since the absence of the chain would reflect a technical problem with Sanger sequencing and not the true biology. Therefore, we used sensitivity as a metric to compare different tools. Firstly, we identified genes of productive HC and LC (K or L) of the ground truth for each cell in each dataset as described in the chapters above. Secondly, we used different tested tools to assemble and annotate HC and LC (K or L) in different experimental libraries that we generated as described above. Then, we compared genes of each productive chain of the ground truth to the corresponding productive ones obtained by a tested method. As IgBLAST and tested tools can sometimes output more than one of each V-D-J germline genes, a HC was defined as a true positive (TP), if at least one of each of the V-D-J genes of a productive HC in the ground truth matched one of each of the corresponding V-D-J genes obtained by the computational method. In case a HC chain was not reported as assembled and productive by the tool, or when at least one of the V-D-J genes in the ground truth did not match any of the corresponding V-D-J genes obtained by a computational method, it was considered a false negative (FN). The same approach was used for the LC, but only V-J genes were matched. Knowing that a B-cell can have two LC and (a process named allelic inclusion, (45)) in case two LC were assembled in the Sanger, we compared only the productive one to the one obtained by the given method. Sensitivity was then calculated using the formula below:

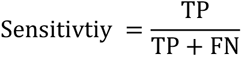

#### Accuracy as a performance metric for simulated synthetic BCRs

In the simulated synthetic dataset described above, we counted both productive and non-productive HC and LC (**Supplementary Figure S2 A** to **D**). In humans, mature antigen-specific B-cells have one productive HC and LC after SHM, as cells that fail to display functional BCRs are negatively selected and undergo apoptosis. However, the introduction of SHMs by immuneSIM returned several HC and LC with out-of-frame V-Js which we interpreted as a proxy of the SHMs in the variable regions, resulting in non-productive chains which would be negatively selected in the real world. Having assumed this, we counted true positive (TP), false negative (FN), false positive (FP) and true negative (TN) chains for each tool to compute the accuracy. Here, TP and FN were calculated the same way as for the sensitivity. If assembled and non-productive chains in the ground truth were marked as assembled and productive by the tools, such chains were called FP. In case highly mutated and non-productive HC and LC in the synthetic ground truth were reported as non-productive by the tool, they were TN. To evaluate this, we computed the accuracy:

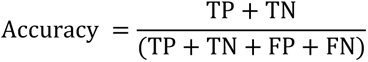

#### Calculation of gene overlap among the methods

To sustain the sensitivity as a metric for performance evaluation of the tested methods, we overlapped the V-(D)-J genes, mapped during the BCR reconstruction by each tool, in a pairwise manner separately for HCs and LCs. To do this, we counted all the HCs that were reconstructed in each dataset by at least one method, and we used this number as denominator of our calculation. This was necessary because some methods were incapable of reconstructing the entire number of HCs in each dataset. Afterwards, we overlapped lists of separate V-(D)-J genes reconstructed by the methods in a pairwise manner and divided this number by the denominator. The same procedure was also performed for the LCs (**Supplementary Table S6**).

#### Assessment of the execution time of each method

We considered the execution time as time needed to reconstruct a BCR from fastq files. For each tool, samples were run as parallelized jobs, in which a sample was run in parallel to all the others at the same time. We first created a multi-node high-performance computing (HPC) cluster employing SLURM (version 18.08.5-2) as a job scheduler with 10 nodes using the Amazon Elastic Compute Cloud (Amazon EC2) and the r5.2xlarge instance that uses the Intel Xeon® Platinum 8000 processor, 3.1 GHz with 8 vCPUs, and 64 GB of RAM. All tools were then run on this virtual cluster requiring at least 2 CPUs and 6 GB of RAM for each job using the following parameters “-n=2” and “--mem-per-cpu=6G“.

#### Final score to evaluate the methods

For each experimental dataset, we aggregated the sensitivity scores of HC and LC separately. For the HC, we summed the sensitivity values across different coverages and read lengths for all the libraries and divided this value by the number of libraries tested (where a library is a dataset with a defined read length and coverage). Given that BASIC was the only method capable of reconstructing HC in libraries of 25 bp length and that BRACER could not reconstruct HC when the read length was 50 bp in most of the libraries, we divided the sum of the sensitivity values across the different libraries of each dataset, by the number of libraries in which the reconstruction was successful. The same was performed for the LC. Afterwards, the two mean values obtained for the HC and the LC were added up and divided by two. This score represents the evaluation of each tool for a given experimental dataset.

For the simulated SHMs dataset, we first summed up the accuracy values of the HC reconstruction across the four SHMs libraries (where each library contained HC with a specific amount of SHMs) and divided this number by the total number of libraries. We then performed the same procedure for the LC. These two values were added up and divided by two to obtain a score for this dataset.

Finally, we computed a cumulative average score by averaging the scores obtained from the four datasets and used it as a final evaluation metric of each method.

#### Creation of the scBCR docker image and data usage

The *scBCR* docker image that was used to install and run the methods tested in this study was built using Debian 10.6 docker. It embeds various tools that share dependencies of different versions that can potentially be incompatible. To avoid this, different Anaconda environments were used to separate different versions of the same software. The native image has BRACER commit 131c2b9, MiXCR 3.0.13 and TRUST4 1.0.4. The conda *omics* environment contains BALDR commit 461d9b0. The conda *vdjpuzzle* environment contains VDJpuzzle 3.0 and BASIC 1.5.0. Along with the files needed to run each tool, the image also contains human genome and annotation version GRCh38, human transcripts, and Bowtie index version GRCh38. The IGMT annotation is also provided to run the tools (46). All the tools were run using standard parameters for all the datasets, if not stated differently. For VDJpuzzle, we had to modify the 1^st^ line of each fastq read when running samples from Upadhyay dataset. Each pair of reads was modified from e.g., “@SRR6471013.1 1” to “@ SRR6471013_1/1” for read 1, and from “@SRR6471013.1 2” to “@ SRR6471013_1/2” for read 2. In case of BALDR, we used the output from the “Ig_mapped & Ig_mapped + Unmapped” model to evaluate the reconstructed HC and LC since it was reported as the best option by the authors (28). In case of BASIC and BRACER, we took the sequences of a given chain and used Change-O (35) within the Immcantation framework v 4.0 (35) to obtain the germline genes, productivity and in-frame V-J junctions. This information was already present in the output folders of TRUST4, VDJpuzzle, BALDR and MiXCR.

#### Validation of reconstructed tetanus toxoid-specific BCRs

We selected three pairs of reconstructed HC and LC belonging to three different BCRs for experimental validation. These tetanus toxoid-specific (TT+) BCRs were part of clonotypes identified in our dataset containing (TT+) memory B-cells. We obtained these clonotypes by running BRACER function “*bracer summarise*” (**Supplementary Figure S4**). We synthesized and cloned variable domains of the assembled HC and LC sequences to produce and test the resulting monoclonal antibodies (mAbs). Firstly, sequences were codon-optimized via GeneArt Gene Synthesis (Life Technologies) and the HC/LC variable genes together with 5’-BamHI and 3’-XhoI restriction sites, the Kozak sequence, and the respective leader sequence were ordered from GeneArt (Life Technologies). The constructs were then ligated into a pcDNA3.1 (+) expression vector (Invitrogen) carrying the IGHG1/4 or the IGLC1/IGKC constant regions (UniProt) respectively, flanking a 3’-XhoI site. The recombinant monoclonal antibodies were produced in Freestyle^TM^ 293-F cells (Gibco) as previously stated (47). Supernatants were harvested 5-6 days post-transfection. IgG antibodies were purified using a 1 ml HiTrap^®^ Protein G HP affinity column (GE Healthcare) followed by a direct buffer exchange using a 53 ml HiPrep^TM^ 26/10 Desalting column (GE Healthcare) according to the manufacturer’s instructions. An IgG enzyme-linked immunosorbent assay (ELISA) was used to determine IgG concentrations of the mAbs according to manufacturer’s protocol (Bethyl laboratories). Anti-TT reactivity was assessed by a TT ELISA. Tetanus toxoid (NIBSC) was directly coated onto C96 Maxisorp NuncImmuno plates (Thermo Fisher Scientific). mAbs were tested in multiple concentrations. Bound anti-TT IgG was detected by polyclonal rabbit anti-human IgG horseradish peroxidase (Dako), ABTS and H2O2 (Sigma-Aldrich).

## RESULTS

### Effect of coverage and read length on BCR assembly, productivity, and sensitivity

To evaluate the experimental parameters affecting the performance of the BCR reconstruction tools in different B-cell types, we generated datasets of distinct library lengths and read coverage. To comprehensively test this, we selected three datasets with different characteristics. The first dataset named “Leiden”, consisted of reads obtained from tetanus-toxoid specific memory B-cells and memory B-cells with unknown antigen specificity. It was generated as part of this study using the SMART-seq2 protocol and had read length of 100 bp. The Canzar (SPEC-seq) (26) and Upadhyay (SMART-seq2) (28) datasets were obtained by sequencing plasmablast cells, yielding read lengths of 50 bp and 100 bp, respectively. We downsampled all three datasets to create libraries with variable coverage levels (from 50,000 to 1,250,000). Similarly, read trimming was done to simulate libraries with different read lengths (from 25 bp to 100 bp). Finally, we assessed whether a set of productive HC and LC (LcK or LcL) could be assembled by each tool for each single cell in the total set of 70 different libraries (see **Material** and **Methods**) and used the ground truth information (see **Supplementary Figure S1 A to C**) to compute the sensitivity (see **Methods**).

### Leiden

We observed different outcomes for the different methods in terms of the percentage of cells with assembled and productive HC and LC (**Figure 2A**) using libraries, generated from this dataset of 72 cells. Specifically, the number of assembled and productive HC was dependent on the number of reads for all the tools and increased with the rising coverage of the libraries. Moreover, this effect was strikingly pronounced for BASIC, which was able to assemble only less than 40% of the HC in libraries with 100,000 reads. Furthermore, BRACER showed difficulties in reconstructing HC in a scenario of 50 bp libraries with a very pronounced effect for coverages below 500,000 reads (**Supplementary Figure S5 A**). BALDR, BRACER and TRUST4 displayed a similar high performance, assembling HC in up to 90% of the single cells. Finally, the average sensitivity for the HC remained relatively stable across the different coverage levels with an average value of 85% for BRACER, followed by BALDR with 77% and TRUST4 with 66% (**Figure 2A** and **Supplementary Table S1**).

**Figure 2.**
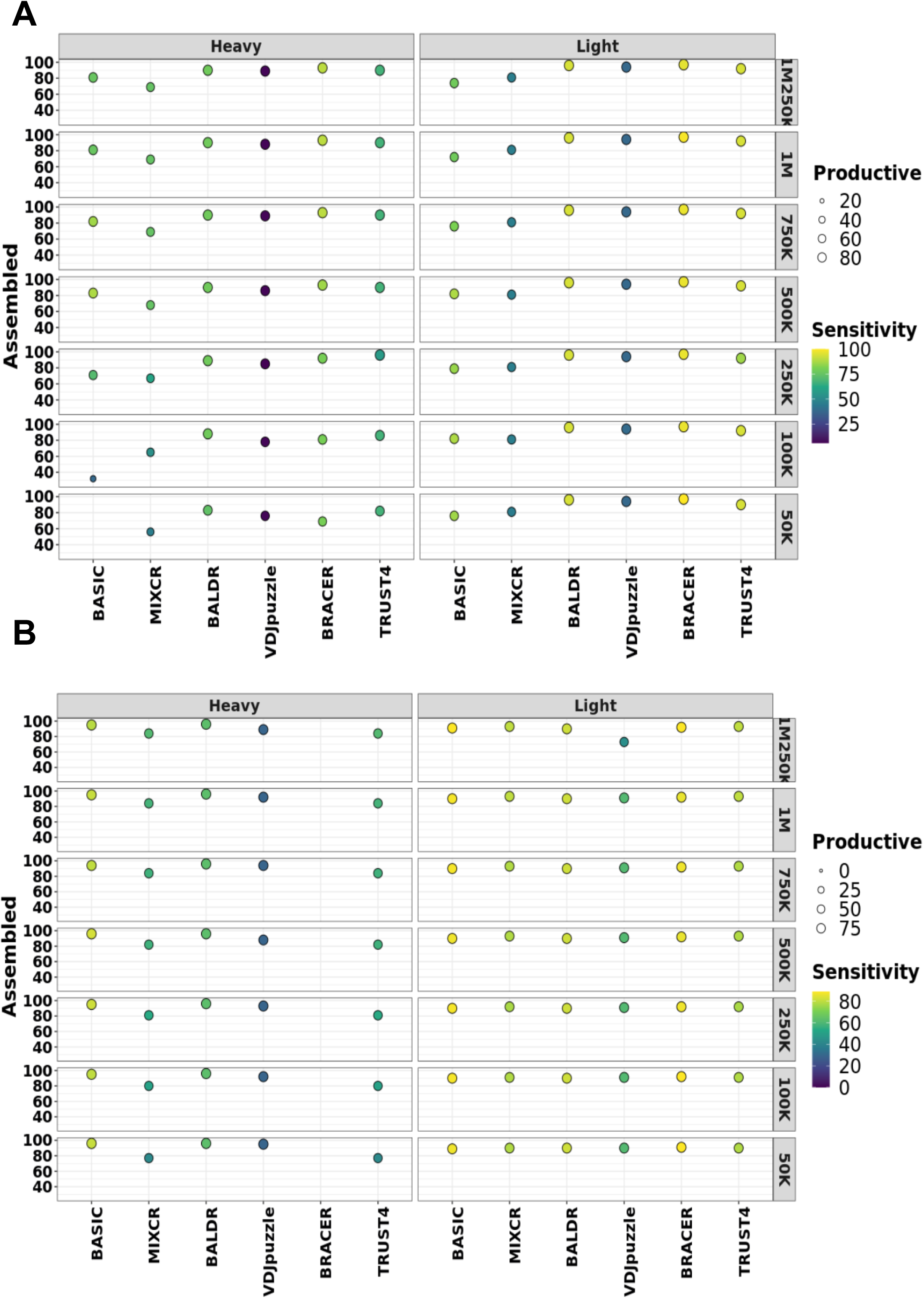
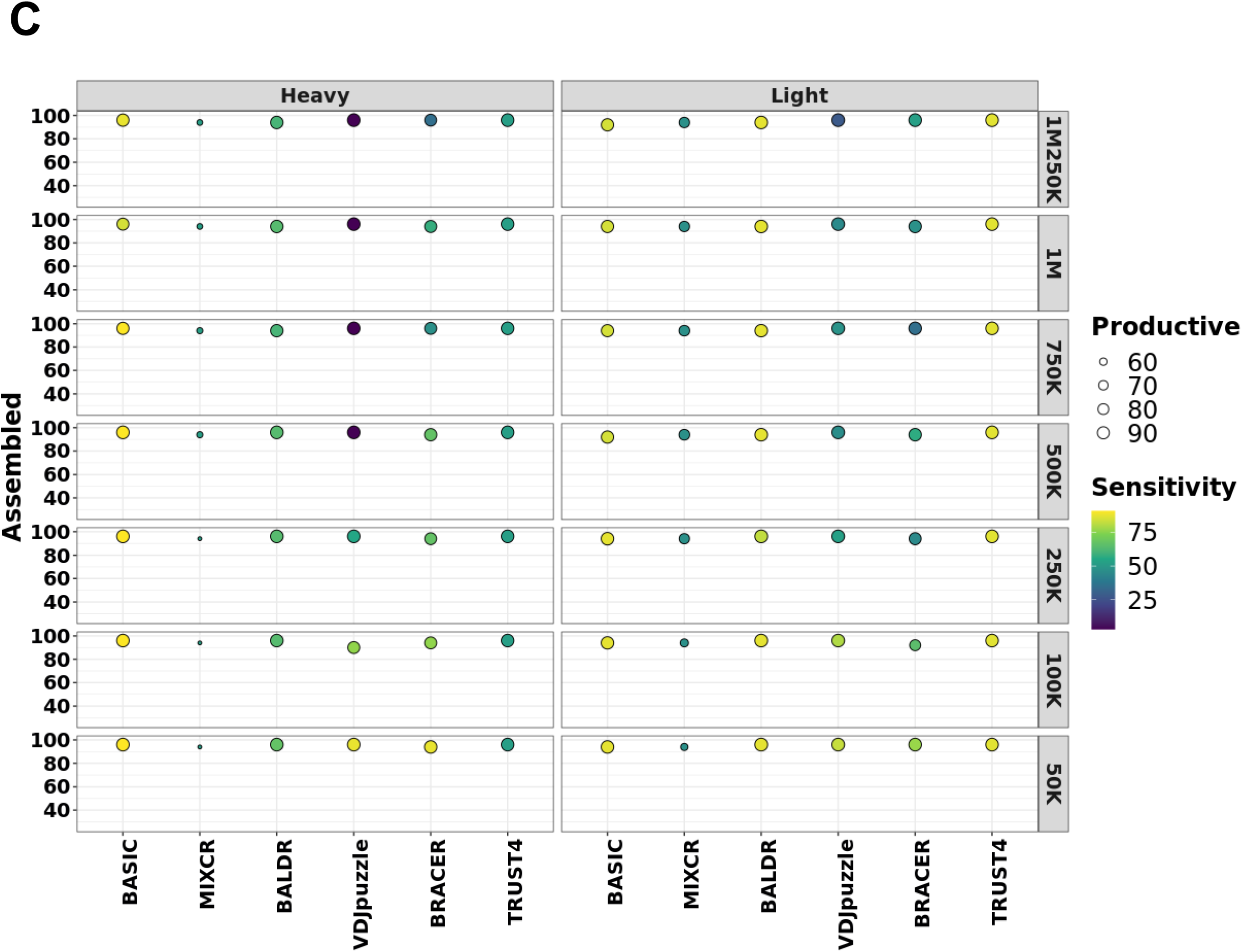
Performance of each method on different datasets. Effect of sequencing depth on the assembly, productivity, and sensitivity of each tool in the **(A)** Leiden dataset created in this work, **(B)** Canzar dataset and **(C)** Upadhyay dataset, consisting of 72, 51 and 113 paired-end single-cell libraries. Original libraries were generated using either SPEC-seq or SMART-seq2 technology and had 100 bp, 50 bp and 100 bp long reads, respectively. These datasets were downsampled to different coverages as displayed in the figures to test the effect of coverage on the assembly, productivity, and sensitivity of BCR heavy and light chains (LcK or LcL). *Assembled* are heavy and light chains without stop codons. *Productive* are assembled heavy and light chains with in-frame V-J junctions. Left y-axis depicts % of assembled chains over the total number of single cells in each dataset. Right y-axis corresponds to coverage. The size of the circles is proportional to the % of the productive chains. Higher intensity of the yellow color of the circles corresponds to the higher sensitivity.

All tools demonstrated a consistently high number of assembled and productive LC (K or L) across the different coverage and read length levels, with BALDR, BRACER and TRUST4 assembling LC in more than 90% of single cells (**Figure 2A**). Likewise, BRACER, BALDR and TRUST4 reached LC assembly sensitivities of 99%, 94% and 93% respectively, being the highest among the tested tools (**Figure 2A** and **Supplementary Table S1**). In conclusion, BRACER, BALDR and TRUST4 were the best performing tools, assembling HC and LC with the highest sensitivity in this dataset.

### Canzar 2017

In contrast to the other two datasets, we could only test the tools using 25 bp and 50 bp read length scenarios due to the nature of the original libraries of 113 plasmablasts in the Canzar dataset. As in the Leiden dataset, we observed varying effects of the tested parameters on the different tools. (**Figure 2B**).

Surprisingly, BRACER failed to assemble HC in all the tested libraries, seemingly due to the shortness of reads. On the other hand, all the other tested tools successfully assembled more than 75% of the productive HC. Moreover, BASIC reached up to 95% sensitivity across different coverage levels for the HC, followed by BALDR and TRUST4 with a sensitivity of 63% and 54% respectively (**Figure 2B** and **Supplementary Table S1**). Although, MiXCR and VDJpuzzle assembled comparably high percentages of the HC, they were less sensitive than others.

Furthermore, all the tested tools successfully assembled LC in more than 80% of the single cells. Additionally, changing coverage didn’t affect the assembly and productivity rates, except for VDJpuzzle, where we observed a drop to 75% of the assembled LC in the 1,250,000 read libraries. Moreover, BASIC and BRACER showed the highest sensitivity of 88% that remained stable across the different coverage levels. Other methods were less sensitive, with BALDR, TRUST4, MiXCR and VDJpuzzle reaching the sensitivities of 82%, 79%, 78% and 59% respectively (**Figure 2B** and **Supplementary Table S1**). Notably, BASIC was the only method capable of assembling productive HC and LC from 25 bp libraries, displaying 54% sensitivity in reconstructing HC, and 88% sensitivity for LC, with the average of 71% for both chains (**Supplementary Figure S6** and **Supplementary Table S1**). Altogether, BASIC, BALDR and TRUST4 were the methods capable to assemble the highest number of productive HC and LC with the highest sensitivity values in the Canzar dataset.

### Upadhyay 2018

Comparably to the Canzar dataset, the Upadhyay dataset was originally generated by sequencing 51 single plasmablasts in a paired-end mode, using, 100 bp libraries instead of 50 bp. This could potentially explain some of the differences we observed (**Figure 2C**). The number of assembled and productive HC across the different coverage levels was equal to or higher than 85% for all tools analyzed except for MiXCR, which assembled less than 60% productive chains. Consistently with the Canzar dataset, BRACER was not able to assemble HC when the length of reads dropped to 50 bp and below (**Supplementary Figure S7A**). Similarly, BASIC achieved the highest average sensitivity value (90%) across the different coverage levels for the HC. It was followed by BALDR and TRUST4 with 64% and 52% of sensitivity, respectively. Finally, VDJpuzzle and BRACER showed an inverse sensitivity-coverage relationship, with both reaching the sensitivity of 88% in libraries made up of 50,000 reads.

Only MiXCR failed to achieve more than 90% of assembled and productive LC chains across the different coverage levels. Among the remaining tools, TRUST4, BALDR and BASIC were the most sensitive, reaching average sensitivity levels equal to 87%, 87% and 86%, respectively (**Supplementary Table S1**). As for the HC, the sensitivity of VDJpuzzle and BRACER increased up to 83% and 77% with the simultaneous decrease of the number of reads to 50,000. An investigation of the read quality of all samples (**Supplementary data 1, Supplementary Figure S3, Supplementary Table S5**) revealed that the Upadhyay libraries harbored an overrepresentation of *k*-mers and duplicated reads which might explain the inverse sensitivity-coverage relationship observed for BRACER and VDJpuzzle. Thus, BASIC was the method capable of assembling the highest number of productive HC and LC with the highest sensitivity value followed by BALDR in the Upadhyay dataset.

### Method sensitivity is reflected by the V-(D)-J gene overlap

Next, we asked whether the performance of the tested methods was driven by the mere availability of a subset of HCs and LCs and the corresponding ground truth sequences in each dataset (see **Supplementary Figure S1 A to C)** or by the capability of each method to detect the same genes and, therefore, alleles during the mapping procedure before reconstruction. To comprehensively test this, we did a pairwise comparison of separate V-(D)-J genes that were reconstructed in HCs and LCs by different methods. We selected libraries of 100 bp with 1.25 million reads from the Leiden and Upadhyay datasets and 50 bp with 1.25 million reads from the Canzar dataset for this analysis (**Figure 3**). Starting with the Leiden dataset (**Figure 3-A**), we observed that BRACER and BALDR had the highest overlap of annotated HC V-(D)-J genes (0.88, 0.72 and 0.84). Moreover, although MiXCR and BRACER had a high overlap of the D genes, a poor overlap of the V and J genes was reflected in a general lower sensitivity of this tool. Furthermore, BALDR and BRACER showed the highest overlap of the V and J genes of LCs (0.9 and 0.9). These results were in line with the overall better performance of these two tools on the Leiden dataset when the ground truth was used. Besides, we observed that BASIC and BALDR had a consistently good overlap of annotated HC V-(D)-J genes (0.71, 0.51 and 0.76) and the best overlap of annotated LC V-J genes (0.86 and 0.8) when compared to the match among other tools while using the Canzar dataset (Figure 3-B). It is important to note that BRACER did not show any overlap of HC genes with any of the tools due to the inability of BRACER to reconstruct HCs using 50 bp read length libraries. Finally, BASIC and BALDR had consistently the best overlap of V-(D)-J genes of the HC (0.86, 0.59 and 0.86) and LC (0.9 and 0.92) in the Upadhyay dataset (**Figure 3-C**). Again, this agreed with the best performance of these two tools on this dataset. Overall, this analysis, which did not rely on the ground truth, reflected the performance of each method which was evaluated when using the ground truth and the sensitivity as a metric.

**Figure 3.**
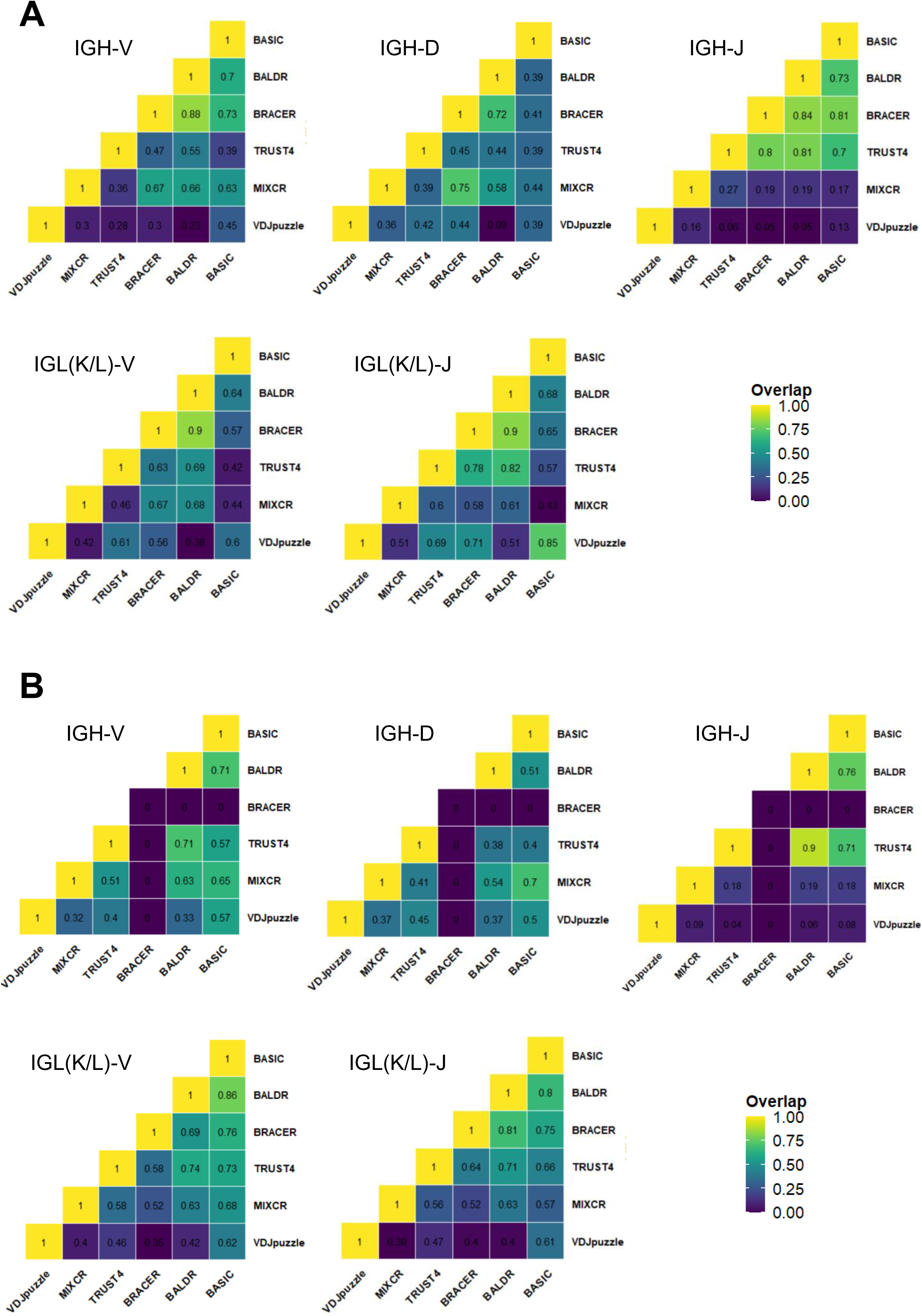
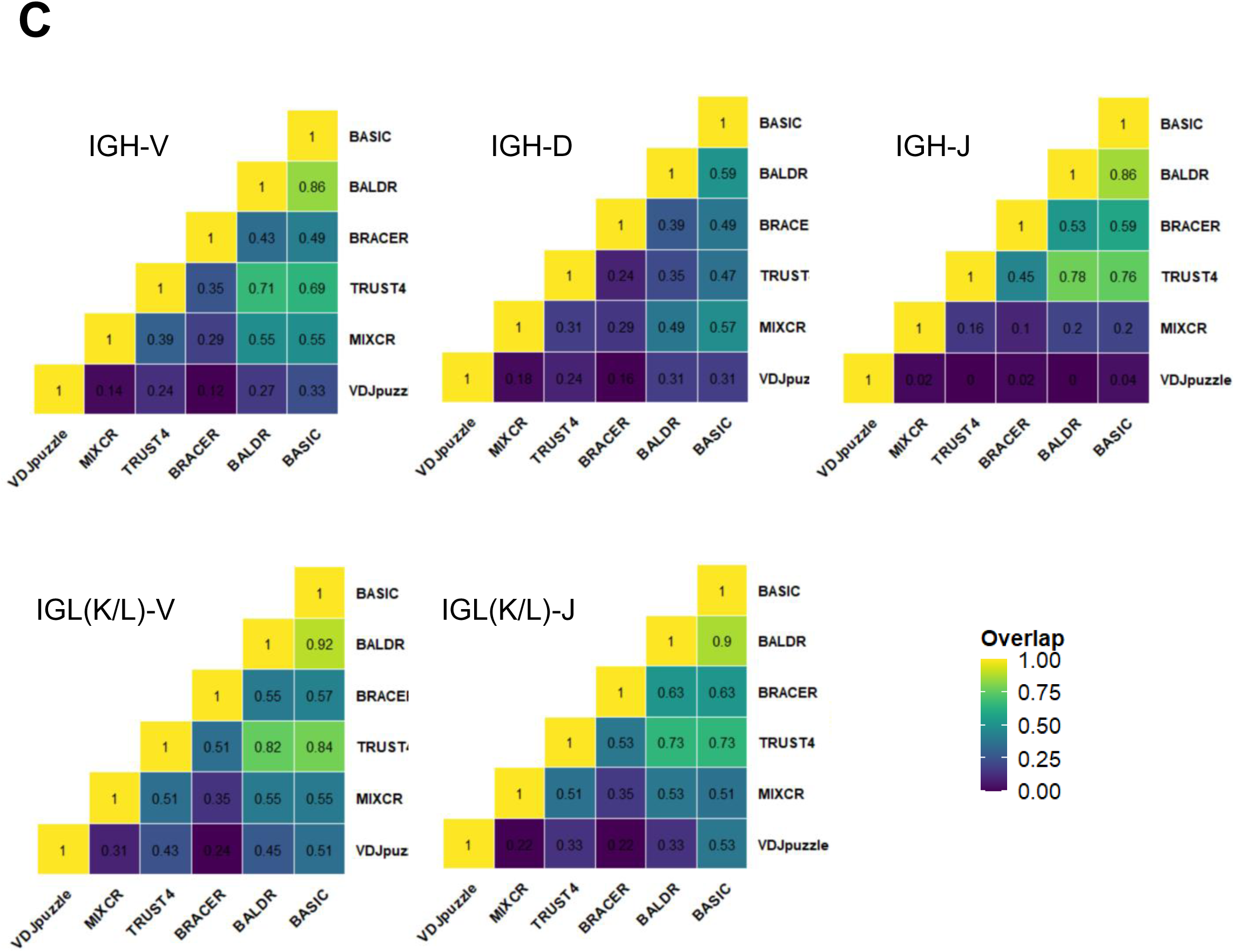
Pairwise overlap of V-(D)-J genes. Proportions of overlapping separate V-(D)-J genes among different computational methods for heavy chains (IGH) and light chains (IGL(K/L)) in the A) Leiden, B) Canzar and C) Upadhyay datasets. Higher intensity of the yellow color in heatmaps corresponds to the higher overlap.

### Effect of somatic hypermutations on BCR assembly, production, and accuracy

After assessing the effect of the read coverage and length on BCR *assembly* and *production*, we next investigated the consequences of different levels of SHMs in the CDRs of the variable domains of the BCRs on the performance of each tool. To address this, we firstly simulated datasets that consisted of 100 HC, 100 LcK and 100 LcL and contained distinct levels of SHMs (ranging from 15 to 60) in the CDRs of their variable domains. Secondly, we created synthetic Illumina libraries and tested the capability of each method to reconstruct HC and LC harboring various levels of SHMs. Finally, we annotated the simulated HC and LC and used them as the ground truth to compute the accuracy of each tool (see **Methods** and **Supplementary Figure S2 A** to **D**).

The first noticeable outcome of this experiment was a pronounced decrease in the percentage of assembled HC and LC (LcK and LcL) with the increasing number of introduced SHMs in the CDRs of the variable domains for MiXCR, VDJpuzzle and TRUST4 (**Figure 4**). Interestingly, we observed a stronger effect of SHMs on the assembly of LC than HC for MiXCR and BALDR compared to other tools. In contrast, BASIC, TRUST4, VDJpuzzle and BRACER did not show large differences in the assembly of HC and LC. Importantly, not all simulated HC and LC were productive in all the datasets (**Supplementary Figure S2 A** to **D)**. In fact, the number of non-productive chains increased with the higher number of SHMs. Consequently, using the tested tools to assemble BCRs from such libraries would result in non-productive HC and LC. Using the proposed metric of accuracy, we found that BRACER returned the highest average values of 95% and 97% across different SHMs levels for HC and LC, respectively (**Figure 4** and **Supplementary Table S1**). This was reflected by the capability of this method to correctly assemble a chain and assign its (non-)productivity as in the ground truth. Other tools, such as BASIC, BALDR and TRUST4, were also stable across the different SHMs levels in HC and LC but demonstrated on average lower accuracy of 66%, 77% and 83% for the HC and 92%, 87% and 79% for the LC, respectively (**Figure 4** and **Supplementary Table S1**). In conclusion, BRACER was the most accurate tool across different SHMs levels in correctly assembling HC and LC for this dataset, followed by BALDR.

**Figure 4.**
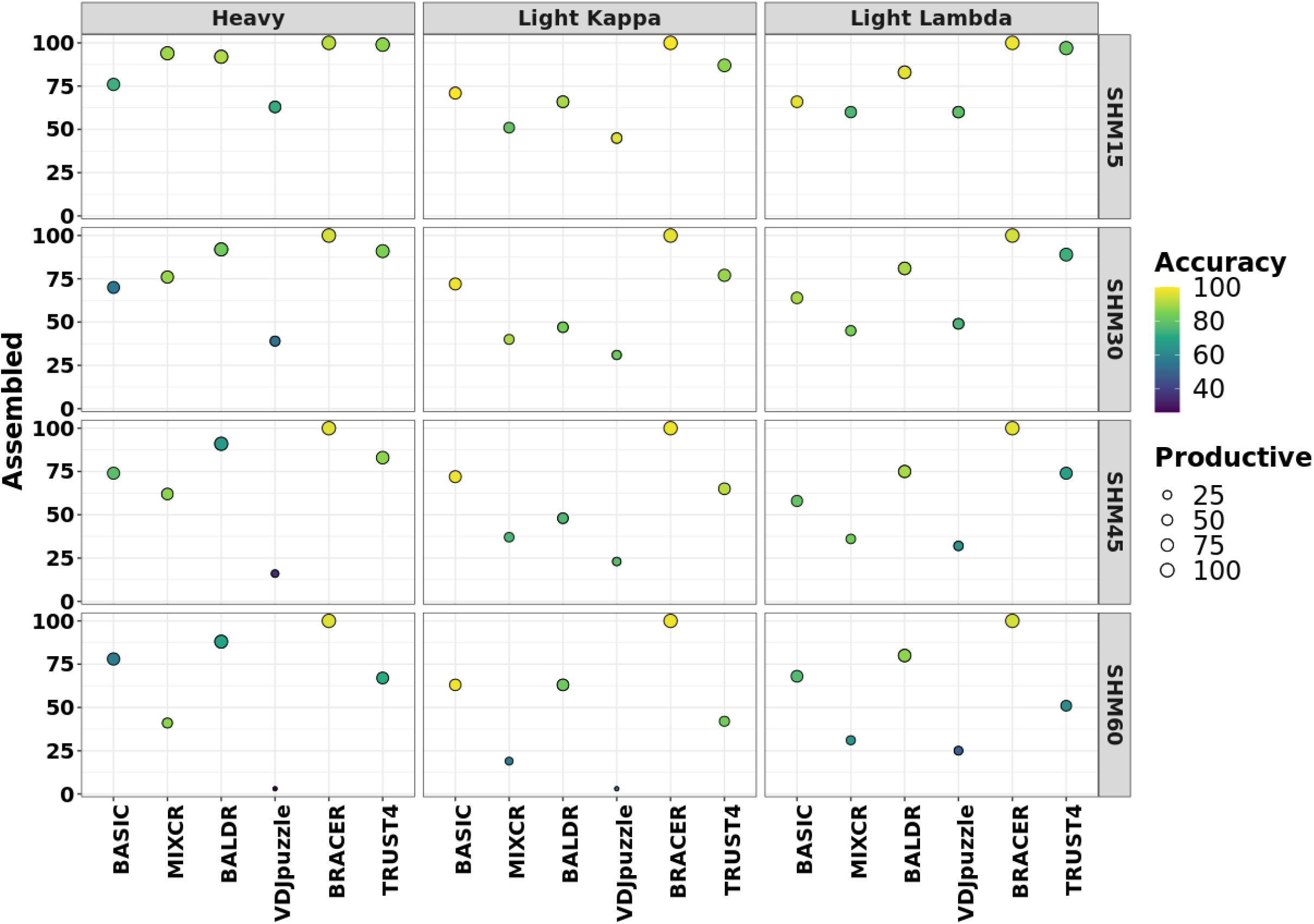
Effect of the number of somatic hypermutations in the variable domain on the performance of each method. 100 heavy (HC) chains, 100 light kappa chains (LcK) and 100 light lambda chains (LcL) were simulated with immuneSim while introducing different amounts of somatic hypermutations (SHMs) in the CDRs of the variable domain (SHMs from 15 to 60). Using these sequences as a reference, Illumina libraries were created with ART (44) tool and all the methods were tested using those libraries. The obtained HC, LcL and LcK were compared to the initially simulated sequences to assess the assembly, productivity, and accuracy rates of each method. *Assembled* are HC and LcK / LcL without stop codons. *Productive* are assembled HC or LcL/LcK chains with in-frame V-J junctions. Left y-axis corresponds to the % of assembled chains. Right y-axis shows the number of somatic hypermutations in each simulated library. The size of the circles is proportional to the % of the productive chains. Higher intensity of the yellow color of the circles corresponds to the higher accuracy.

### Validation of the specificity of assembled BCRs

To validate the specificity of the BCR sequences assembled by the different algorithms, we produced three mAbs using productive BCR sequences that were assigned to three different clonotypes by BRACER using the SMART-seq2 Leiden dataset obtained in this study. Two mAbs were based on BCR sequences of single tetanus toxoid (TT)-specific B-cells collected from patient 1 (cell B10 and cell D8) and a third mAb was based on a sequence isolated from patient 2 (cell G1) (**Figure 5A** and **Supplementary Figure S4**). Firstly, the BCR sequences of these three cells assembled by all algorithms were compared to the ground truth that was obtained using ARTISAN-PCR followed by Sanger sequencing. The assembled sequences of 1-B10 LC, 1-D8 LC and the 2-G1 HC were identical among all different algorithms and the ground truth. However, MiXCR assembled two differences when compared to all other algorithms and the ground truth in the sequence of 1-B10 HC. In the 1-D8 HC and 2-G1 LC, several differences between the ground truth and the output of all algorithms were found (**Figure 5B**). We decided to produce the mAbs based on the sequences assembled by BRACER because two of the three selected sequences showed clonal relationship with BCR sequences determined by this tool in the dataset of TT-specific B-cells (**Figure 5C**). After cloning and expression of selected mAbs, an IgG ELISA was first performed to stratify concentrations up to 1.5 µg/ml followed by a TT ELISA to test the tetanus toxoid binding of these antibodies. We confirmed that an unrelated mAb with a proven specificity for citrullinated antigens was negative in the TT ELISA, while a validated TT mAb was positive. All three tested mAbs showed a clear TT binding, with 1-D8 displaying the strongest signal (**Figure 5D**). In conclusion, except for sequence regions in which all the tools reported a different nucleotide composition when compared to the ground truth among themselves, only MiXCR reported individual amino acid differences in the CDR2 region. Nevertheless, the experimentally validated clonotype-forming BCRs showed high antigen specificity.

**Figure 5.**
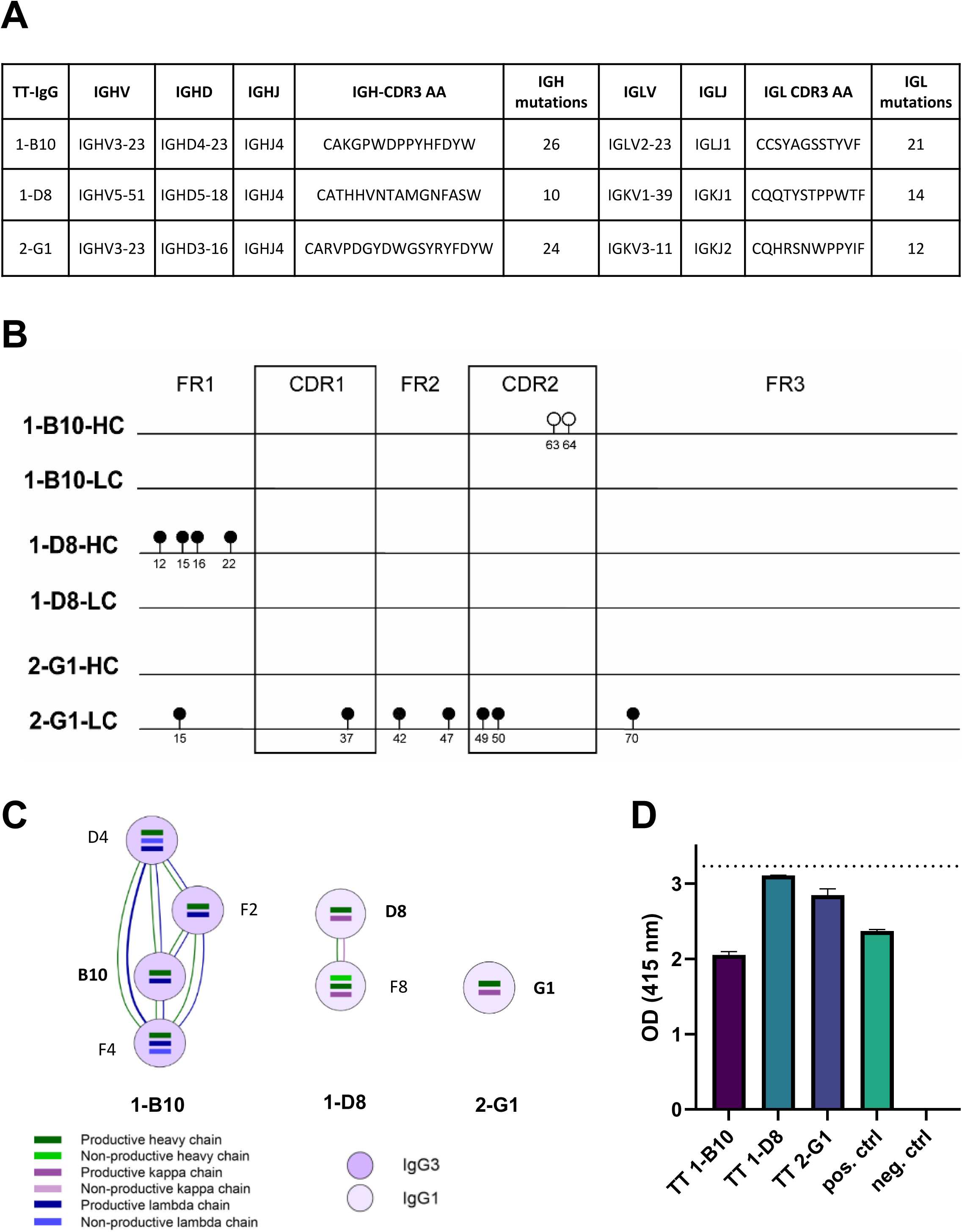
Validation of the tetanus-toxoid specificity of the monoclonal antibodies that were produced based on the assembled BCR sequences. **(A)** Characteristics of the tetanus toxoid (TT)- IgG BCR variable region of the selected mAbs, based on the full-length BCR sequences obtained from the ground truth. V-D-J, CDR3 amino acid (AA) sequences and the numbers of nucleotide (nt) mutations compared to the germline sequence are depicted for the heavy (HC) and the light (LC) chain of each TT-IgG monoclonal antibody. **(B)** Sequence alignment of the HC and LC chains of the sequences used for mAb production. The lollipops depict differences between the ground truth and the used tools. The positions of the lollipops are based on the IMGT amino acid numbering. Open circles depict differences of the MiXCR assembled sequence compared to the ground truth and the output of all other tools. Closed circles depict differences between the ground truth and all the evaluated tools. **(C)** The clonotype networks of the selected BCRs with other TT+ single cells obtained using BRACER (27). **(D)** Antibody validation using a tetanus toxoid ELISA for all antibodies. The ELISA was performed with a concentration of 1.5 µg/ml of the mAbs. Validated mAbs with (pos. ctrl) and without (neg. ctrl) tetanus-specificity were used as controls. The optical density (OD) was measured at 415 nm 20 minutes after ABTS addition. The dashed line shows the upper detection limit of the TT ELISA.

### Evaluation of the execution time

If an experiment includes thousands or millions of cells to be assembled, the runtime during which tools complete BCR assembly might become a determining factor. Therefore, we evaluated the execution time of each tested method using a standard virtual cluster (see **Methods** and **Supplementary Table S2**). As a result, we observed that runtimes increased with the increase of coverage and read length for all methods, except BRACER that demonstrated stable execution times across all coverages, starting from 50 bp (**Figure 6** and **Supplementary Figure S8 A** to **D**). Finally, TRUST4 was the tool that processed the highest number of reads per second, followed by MiXCR, BASIC, BALDR, BRACER and VDJpuzzle.

**Figure 6.**
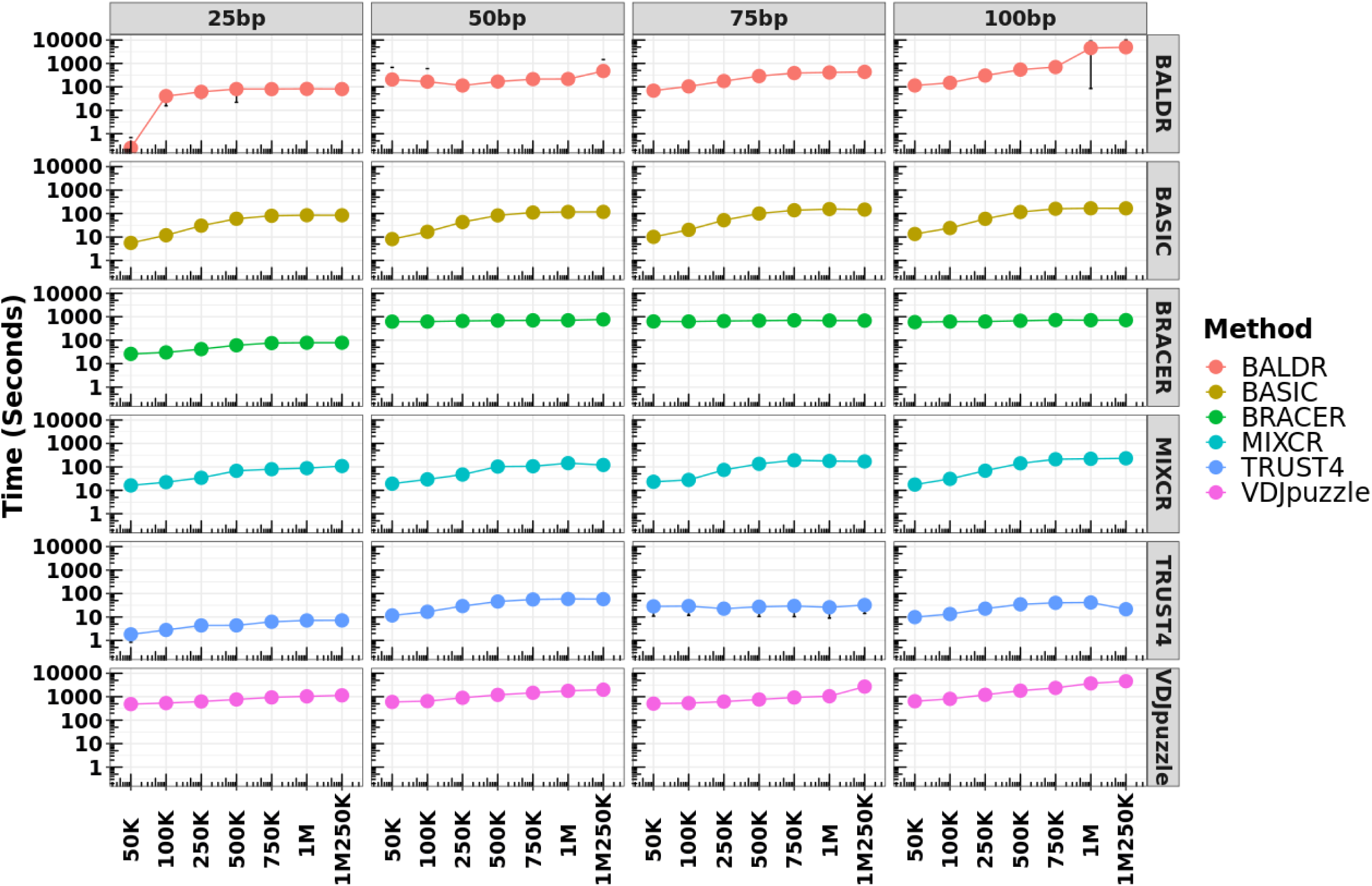
Execution time of each method using samples with different read length and coverage. Each method was tested using standard parameters and libraries with different read length and coverage. These libraries were simulated using the Leiden dataset. Each dot represents the median value (seconds) of the execution time calculated using all single cells from a particular library type. Top x-axis depicts the read length of the libraries. Bottom x-axis shows the coverage of the same libraries. Left y-axis corresponds to the runtime in seconds. Right y-axis shows the tested tool. The circle of each measurement corresponds to the median value of the time of execution of all the cells for a particular library type. The deviation from this value, represents the standard deviation.

## DISCUSSION

In this work, we extensively benchmarked six computational methods for BCR reconstruction using four different B-cell datasets, three experimental and one simulated. We focused our attention on the evaluation of methods capable of reconstructing full-length BCRs. The primary aim of this work was to provide guidance for the method of choice for different plate-based scRNA-seq datasets and scenarios (**Figure 7C**). In addition, we directed our attention to understanding the performance of each method on highly mutated BCRs which are common in autoimmune diseases (7, 10), cancers (6, 11) and in neutralizing anti-HIV antibodies (32–34).

**Figure 7.**
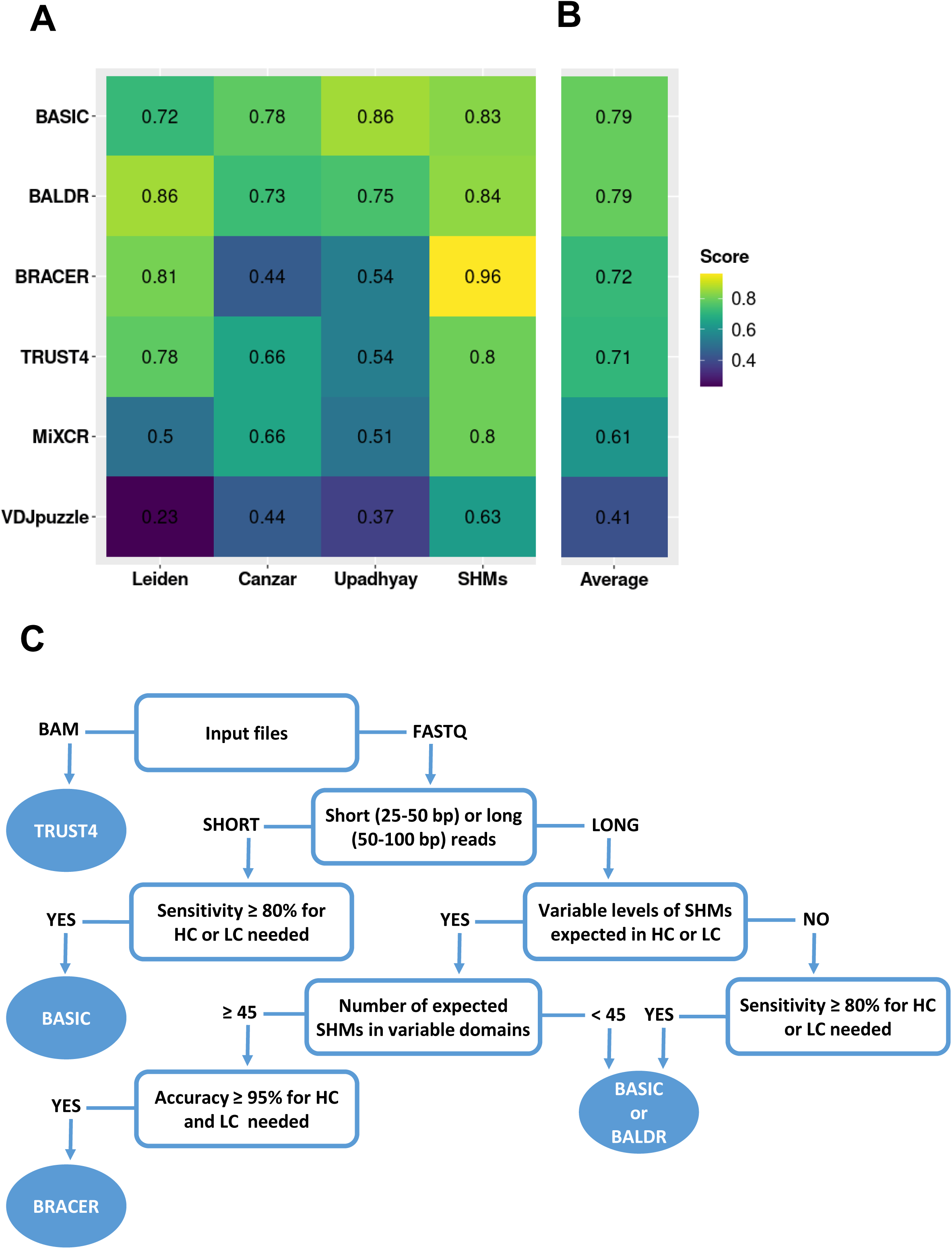
Average performance of benchmarked BCR methods on the different datasets. **(A)** For each method, a weighted average value for the sensitivity was calculated by averaging single sensitivity values, that were obtained for HC and LC after running the tools on the different libraries of the three datasets: Leiden, Canzar and Upadhyay. A similar procedure was performed using the accuracy for the HC and LC obtained using the simulated SHMs dataset. **(B)** A cumulative average score for each method across the different datasets was obtained. **(C)** Our recommendation tree for method selection, according to the research question and type of the dataset to be analyzed.

Methods based on either “semi *de novo”* (BASIC) or “*de novo”* (BALDR) assembly and using the IMGT annotation during the mapping procedure, showed on average the highest performance when assembling both HC and LC (K or L) of BCRs (**Figure 7A)** in the two plasmablast datasets (Canzar and Upadhyay). Moreover, despite the Upadhyay dataset being biased towards the overrepresented *k*-mers, both BALDR and BASIC algorithms maintained a good overall performance as previously reported (28), with BASIC being independent of the type of immunoglobulin chain. Despite having a similar high performance when compared to the other methods in the Leiden dataset, BASIC was, however, sensitive to the decreasing coverage level when reconstructing HC, and was outperformed by the *de novo* assembly-based methods BALDR, BRACER and TRUST4 (**Figure 7A)**. The main difference of the Leiden dataset compared to the previously published datasets was the type of cells used for library construction, memory B-cells and plasmablasts, respectively. This could have had a potential impact on the results we observed as plasmablasts produce high amounts of antibodies and thus have higher amounts of BCR mRNA. However, an excess of reads which belong to the V-(D)-J genes in a sequencing library does not strictly implicate better performance of all the tools. This could be observed in the Figure 7A for the two plasmablast datasets Canzar and Upadhyay, in the which the final score of all the tools was lower than the one obtained for the Leiden dataset of memory B-cells. This could be explained by an open problem in genomics, which describes a possibility of multiple reads, belonging to a short variable genomic region (in this case V-D-J genes), to map to multiple locations. Moreover, in accordance to the previous study (48), MiXCR showed a generally low sensitivity in respect to the other tools, which was reflected by the inconsistency of genes annotated in the assembled HC and LC, which is probably due to the high number of gene mis-hits. Thus, our results suggest the adoption of BASIC, BALDR and BRACER for the investigation of B-cell repertoires where BCRs are carrying a particular antigen specificity to accelerate antibody-based drug design.

When assessing the capability of each method to reconstruct BCRs bearing variable levels of SHMs in their variable domains, we noticed the limited performance of VDJpuzzle. We propose that this was due to the intrinsic property of the tool to completely discard reads in case they do not map to any of the V-D-J genes and constant regions during the first assembly step (29). To support this notion, methods, based on more sophisticated algorithms to reconstruct the variable portion of the HC and LC by overlapping the unmapped reads to those mapping to V-J junctions together with genes present in the constant regions of HC and LC, showed to have adopted a better strategy. Therefore, BRACER, BALDR, BASIC and TRUST4, should be chosen when assembling highly mutated sequences.

The experimental and simulated datasets used for this benchmarking study supplemented each other. On one hand, the experimental data reflected the real-world scenario of a limited number of available (antigen-specific) B-cells and underlined the challenges in the assembly of their BCRs at different cell differentiation stages. On the other hand, larger sets of sequences with specific characteristics like SHM load with absent sequencing challenges (e.g., duplicated reads and overrepresented *k*-mers etc.) can be compared when using simulated datasets. However, in general, simulated datasets cannot completely reflect and thus replace the *in vivo* situation.

In terms of execution time, TRUST4, MiXCR and BASIC can process the highest number of reads per second. They can run on a machine with as few as 2 CPUs Intel Xeon® Platinum 8000 with 3.1 GHz frequency each and 6 GB of RAM and process single-cell libraries of one million reads in approximately two minutes thus making them adoptable in every modern lab.

Importantly, some of the BCR assembly tools, namely, MiXCR, VDJpuzzle and TRUST4 can also be used to assemble T-cell receptors (TCR) from single-cell sequencing data. However, since BCR reconstruction is complicated by somatic hypermutation, while this is not the case for TCRs, an independent benchmarking of the tools should be performed for TCR assembly.

With the goal to validate the specificity of the BCR sequences assembled by the different algorithms, three monoclonal antibodies were produced. Remarkably, all tools showed consistent sequence (dis)similarities when compared to the ground truth, except for MiXCR, which assembled 1-B10 HC with two additional mutations. This is in line with the previous study (48) showing a high frequency of gene mishits during the alignment step of MiXCR and the resulting negative impact on the BCR sequence assembly. The disagreement of all the tools with the ground truth for the 1-B10 HC was in line with the low quality of the Sanger sequencing in the region where different nucleotides were called, suggesting that the computational methods can be superior to Sanger in such cases. However, the Sanger sequence of the 1-D8 LC was of high quality and may hint to either the amplification bias during the ARTISAN PCR or to the issues during the BCR reconstruction step. Since all computational tools employ different methodologies for BCR assembly, the latter seems unlikely. Finally, the antigen affinity assay showed a clear tetanus toxoid binding by the cloned antibodies. This confirms that the adoption of these tools in the research and clinical setting would be beneficial for BCR and antibody reconstruction.

We would also like to emphasize that this benchmark study was explicitly done with datasets generated using plate-based scRNA-seq techniques, such as SMART-seq2 or a variant of it named SPEC-seq. TRUST4 was the only method, among the publicly available ones, capable of processing 10X BCR data (30). However, the unfeasibility to obtain the ground truth for 10X data and the scarcity of tools to process them made us exclude this type of data from our benchmark. Nevertheless, such comparisons should be addressed in the future studies. Moreover, although BRAPeS (49) was initially included in our evaluation, the extremely long run times to reconstruct just the CDR3 region, resulted in a need to subset the reads (∼5000 as suggested by the developer). Since the scope of this work included understanding the effect of read coverage and length on tool performance, we excluded BRAPeS from further evaluations.

In conclusion, we provide clear guidance to select the best method (**Figure 7C**) according to the data type and research question the user has at the start of the BCR reconstruction experiment to facilitate the research. In our opinion, this work will help to improve the existing and develop new methods for BCR construction, especially adapting them to other sequencing technologies that are gaining increasing popularity, such as those using 10X, Oxford Nanopore and PacBio sequencing platforms.

## AVAILABILITY OF THE DATA AND RESULTS

Figure 2, 4 and 7 can be reproduced using data in the **Supplementary Table S1**. Figure 3 can be reproduced using the data in the **Supplementary Table S6**. Figure 5 can be reproduced using the **Supplementary Table S2**. The Sanger sequencing results of the “Leiden” dataset together with the annotation can be found in the **Supplementary Table S3.** The fasta sequences of the simulated datasets with different levels of SHMs in the CDRs of the variable domains of HC and LC can be found in the **Supplementary Table S4**.

We deposited the *scBCR* docker image together with the instructions to run all the tested methods in the following GitLab repository: https://gitlab.com/tAndreani/scBCR. In addition, we also created notebooks to compute the sensitivity and accuracy as well as to re-create the plots presented in this manuscript within the same directory.

## ACCESSION NUMBERS

The single-cell RNA-seq data provided in this work are available at the SRA, Bio project ID PRJNA783770 (access link until release: https://dataview.ncbi.nlm.nih.gov/object/PRJNA783770?reviewer=4roc8i2si3hfao2q7lv4hovaec). Supplementary Data are available online.

## ACKNOWLEDGEMENT

We would like to acknowledge the Sanofi Global Postdoctoral Fellowship Program for financing the postdoc position of Tommaso Andreani. We thank the Flow cytometry Core Facility (FCF) of Leiden University Medical Center (LUMC), the Netherlands, for the assistance with the cell sorting. We also thank Nima Nouri from Sanofi Precision Immunology group, Franck Rapaport and Taylor Sorenson from Sanofi’s AI & Deep Analytics, Omics Data Science and In Silico Drug Design groups, respectively, for reading the manuscript and providing feedback.

## SUPPLEMENTARY INFORMATION

### B-cell receptor reconstruction tools

We chose algorithms according to the capability to (i) reconstruct full-length B-cell receptors (BCR) consisting of CDR(s) and FWR(s) regions of paired heavy and light chain in a single cell, (ii) process different SMART-seq libraries and where (iii) full documentation was provided to install and run the software.

### MiXCR

The algorithm of MiXCR is designed to process single or paired-end reads from single and bulk RNA-seq data to quantify clonotypes. It performs three main operations: (i) alignment of the reads to V, D, J and C genes of B-cells with a proprietary aligner using IMGT annotation (1), (ii) clonotype assembly using the alignment from the previous step and (iii) export of the alignment results to allow the user to investigate sequence information. The alignment of the reads to V, J, and C genes is based on a specialized version of the *k*-mer chaining algorithm previously proposed by Liao (2), modified to handle short alignments and to determine boundaries of alignments with high accuracy. During the alignment, two main steps occur: in the first step named “seed and vote”, a set of seeds is created using reference sequences. During the voting step, the algorithm performs a heuristic search to maximize a scoring function, calculated based on the number of matching and absent seeds in a certain region of query sequence (sequencing read). Candidate alignments are built for spaces between seeds and for regions outside boundary seeds. In the second step, alignments are built using the classical Needleman– Wunsch algorithm with a banded matrix for sequence between seeds and using a modified Smith-Waterman algorithm for boundary sequences.

After alignment, the algorithm performs clonotype assembly. The assembler tries to extract gene feature sequences from aligned reads (called clonal sequences) specified by the user (by default CDR3) and the clonotypes are assembled with respect to the clonal sequence. If the aligned read does not contain a clonal sequence (e.g., CDR3 region), it is be dropped. Records with clonal sequence containing only high-quality nucleotides are used to build core clonotypes by grouping records with equal clonal sequences (e.g., CDR3). After the core clonotypes are built, MiXCR implements several steps to rescue information from low quality reads that can map to already assembled clonotypes. In case of several matching clonotypes, a single clonotype is randomly chosen with weights equal to clonotype abundances. If no matches are found, the record is finally dropped. After clonotypes are assembled by initial assembler and mapper, MiXCR proceeds to clustering. The clustering algorithm tries to find fuzzy matches between clonotypes and organizes matched clonotypes into hierarchical trees (cluster), where each child layer is highly like its parent but has significantly smaller abundance. After clusters are built, only their heads are considered as final clones.

Finally, clonal sequences are aligned to reference V, D, J and C genes. Since gene features used to build clones (e.g., CDR3) are different from those used in aligner (e.g., V-region, J-region etc.), it is necessary to rebuild alignments for clonal sequences. Since all hits are known in advance, these alignments are built using the slower but more accurate Smith-Waterman algorithm (3).

### BASIC

BCR receptor for single cells (BASIC) is a semi *de novo* assembly method that can reconstruct BCRs from single or paired-end fastq files. The semi *de novo* assembly is performed in two steps: (i) first BASIC uses known constant and variable regions to identify anchor sequences and then (ii) uses these anchor sequences to guide the *de novo* assembly of the BCR. In the first step, BASIC uses a pre-compiled database of the known germline variable and constant regions of the heavy and light chains extracted from the human IMGT database (1). The reads are aligned to each variable and constant region of the HC or the LC, and short sequence windows with the highest number of mapped reads are identified. These regions will be used as an anchor for the next step. In the second step, BASIC performs *de novo* assembly to join the anchor sequences. While doing this, BASIC extends each anchor iteratively in the 3’ direction using a greedy minimum entropy approach. In each iteration, starting with the anchor sequence, all overlapping reads are identified for a given direction. In contrast to other assembly methods that use a fixed *k*-mer size, BASIC conceptually explores a graph structure that connects nodes by an edge if the corresponding read sequences overlap by any length (4). Finally, each anchor is extended in the 5’ direction in the same way. Fully reconstructed highest scoring heavy and light chains are outputted in the fasta format.

### BRACER

BRACER is a pipeline for the *de novo* B-cell receptor reconstruction and clonality analysis. It takes single or paired-end single-cell RNA-seq data as an input, trims the reads at low quality positions and discards the PCR duplicates. During the mapping step, BRACER uses the “combinatorial recombinome” annotation file. This file consists of all possible V-J gene segment combinations present in the IMGT reference (1). Eight ambiguous Ns (N = any nucleotide) are introduced between the V and J gene segments to mask the D region in order to allow the mapping of BCR sequences with a range of different CDR3 regions. In addition, 20 Ns are added at the 5′ end of each recombinant sequence to enable the alignment of reads running into the leader sequence. The mapping is performed using Bowtie 2 (5) with low penalties for the introduction of gaps and alignment against ambiguous N nucleotides in the reference sequence. If the library consists of short 50 bp reads and given that the CDR3 regions of heavy chains can be relatively long, a second round of alignment is performed to extract reads mapping mainly or solely to the CDR3 region, which could not be extracted in the first round of alignment. The second alignment is performed with high mismatch and gap penalties to extract reads that fully or partially overlap with reads from the first alignment step, facilitating the extraction of the CDR3-derived reads. For each locus, reads aligning to the combinatorial recombinome are provided as an input to the RNA-seq *de novo* transcriptome assembler Trinity (6).

As BCRs undergo somatic hypermutation and unknown polymorphisms could exist in BCR gene segments, BRACER does not convert reconstructed sequences to full-length sequences using the assigned V and J sequences from IMGT. Instead, BraCeR runs IgBLAST (7) with IMGT-gapped reference sequences and parses the output files using the Change-O tool suite to extract the CDR3 and junction-sequences and to determine if the contig is productively rearranged. If Change-O fails to parse the IgBLAST output for a contig, the productivity and extraction of the CDR3 sequence is performed using a custom script. In such cases, it is assessed whether the assembled sequences have a CDR3 region in the correct reading frame and whether the sequences lack stop codons. If this is the case, the sequences are classified as productive. For the assessment of in-frame junctions and identification of CDR3 amino acid and nucleotide sequences, the productive recombinants are translated starting from the six nucleotides before the CDR3 start position denoted by IgBLAST (7). CDR3 is defined as the region flanked by the final cysteine residue of the V gene and the conserved WGXG (HC) or FGXG (LC) motif in the J gene. If the sequences lack the conserved [FW]GXG motif in the correct reading frame, the scripts are implemented to search for alternative XGXG, WSQG and FSDG motifs. If none of these motifs are found in the correct reading frame, the scripts are implemented to search for the motifs in the two remaining reading frames in order to extract out-of-frame CDR3 sequences (8).

Finally, after assigning a BCR sequence to each single cell within a dataset, BraCeR performs clonal assignment and network reconstruction of the single cells. For this, productive BCR sequences for each locus across a cell population are clustered into clonal groups using the Change-O toolkit ‘bygroup’ subcommand ‘DefineClones’. As proxy for the clustering, BraCeR uses common V and J genes in the sets of potential V and J genes between the sequences, equal CDR3 length, and CDR3 nucleotide distance < 0.2 calculated using a human 5-mer targeting model, mouse 5-mer targeting model or nucleotide Hamming distance for any other species, normalized by length. After assigning the cells into clusters, a clonal network is built. To visualize the clone groups, BraCer performs network reconstruction using custom scripts. The network is constituted by the following elements: each single cell is represented by a node in the graph, and edges between the nodes represent clonally related BCR sequences. Cells are investigated whether they share a heavy chain belonging to the same heavy chain clone group. In the positive case, the algorithm searches for shared light chains belonging to the same light chain clone group (8).

### BALDR

BCR Assignment of Lineage using *De novo* Reconstruction (BALDR) is an algorithm for the *de novo* assembly of B-cell receptors. As input, it takes single or paired-end fastq reads that are subsequently trimmed for adapters. At this step, the method allows two different modalities in order to perform HC and LC reconstruction: i) using the option “Unfiltered”, all the reads that were trimmed in the first step are directly used to perform *de novo* assembly using Trinity (6); ii) using the second option “Filtered”, reads are mapped using different modalities: a) by mapping them to the IMGT annotated V-D-J genes, b) by mapping them to the Ig loci in human reference genome (GRCh38) and reusing the unmapped reads or c) by using IGH and IGL in the “Combinatorial Recombinome” provided by BRACER (8). Afterwards, the mapped reads are used to run Trinity and to perform *de novo* assembly (6). The IGH and IGL sequences assembled from the “Unfiltered” or “Filtered” modality (depending on the user choice) are annotated using IgBLAST (7) to call the V-D-J genes and predict the CDR3. After the annotation step, the best sequence is assigned to the HC and LC choosing the genes with the highest coverage, while redundant and unproductive chains are discarded. Fully reconstructed, assembled, and productive sequences for HC and LC are outputted together with the gene annotation.

### VDJPuzzle

This algorithm is used to *de novo* reconstruct full-length BCRs using single-cell RNA-seq data with paired-end reads. The workflow starts by trimming the reads and aligning them to the reference genome. Afterwards, reads with at least one read aligned to any of the V, D, J, or constant-region genes are extracted. This step is repeated for each chain. Extracted reads are *de novo* assembled using Trinity (6) and this step is repeated for each chain. The resulting contigs are matched to the IMGT database (1) using MiGMAP (https://github.com/mikessh/migmap) to find complete, productive, and in-frame BCRs. At this step, the primary BCR repertoire for each chain is generated using the contigs that match a BCR chain. Next, reads are re-aligned against the primary repertoire for each chain and the resulting aligned reads are used to generate a second round of contigs. The assembled reads are used as an input for MiGMAP to find complete, productive and in-frame BCRs. Contigs corresponding to BCRs from the primary and secondary repertoires are merged to form the tertiary repertoire and an error correction module is applied to identify single nucleotide mutations occurring due to technical errors of the *de novo* assembly. Finally, reads are realigned to the tertiary repertoire and, if a single nucleotide substitution is found within the tertiary BCR sequence with a frequency of >50%, the dominant nucleotide is chosen as the correct base.

### TRUST4

This algorithm can perform BCR reconstruction from SMART-seq2 and 10x libraries. It has a two-step approach that is followed by annotation. The first step is named “candidate read extraction”, where reads within the fastq or bam files are extracted. The second step is named “*de novo* assembly”. For the annotation step, TRUST4 uses the IMGT database (1).

During the first step, using an alignment file in BAM format as an input, the algorithm searches for a read or its mate, which aligns to V, J or C locus and adds this read to a candidate read set. If a read is unmapped and is not a candidate based on mate information, TRUST4 tests the significance of the read overlap with V, J, or C genes. If there is a significant overlap, this read, and its mate are candidate reads. When the input is based on raw sequence files (e.g., fastq files of SMART-seq2 libraries), TRUST4 applies the significant overlap criterion. To identify whether a read has a significant overlap with one of the V, J or C genes, TRUST4 first locates the receptor gene with the highest number of *k*-mer hits (default=9) from the reads and subsequently computes the longest chain from those *k*-mers to filter incompatible hits. If the data has barcode information, such as 10x Genomics scRNA-seq data, TRUST4 corrects the barcode.

After the candidate read extraction, selected reads are used to perform *de novo* assembly using the “read overlap scheme”. TRUST4 implements a greedy extension approach by aligning candidate reads to existing contigs, one by one. To perform the alignment, TRUST4 builds an index for all *k*-mers in these contigs. Based on overlaps, TRUST4 updates contigs with the following rules: (i) if a read partially overlaps one contig, TRUST4 extends this contig; (ii) if a read partially overlaps several contigs, TRUST4 merges corresponding contigs; and (iii) if a read does not overlap any existing contigs, TRUST4 creates a new contig with this read’s sequence. The most frequent *k*-mers are used as the starting point in the de Bruijn-graph-based transcriptome assembler, Trinity (5). TRUST4 clusters reads with somatic hypermutations into the same contig by representing a contig as the consensus of assembled reads. The weights are considered during read alignment to tolerate the somatic hypermutations in BCRs. For example, if for a particular position nucleotides A and T have high weights on the contig, it is a match if the read has nucleotide A or T. Therefore, reads with different somatic hypermutations can align to the same contig, which avoids the creation of redundant contigs. In the first round of contig assembly, due to read sorting and greedy extension, a contig for the abundant recombined gene attracts all reads from the same V, J and C genes even though these reads come from different recombinations. The mate-pair information fixes this issue by reassigning reads to the appropriate contigs. When input data are SMART-seq, since there is no need to perfect assemblies for low-abundant sequences in a cell, TRUST4 can skip the extension to reduce running time (9).

After reads extraction and alignment followed by the *de novo* assembly, TRUST4 performs the annotation step. Here, TRUST4 aligns the assembled contigs to sequences in the IMGT database to identify V, J and C genes. Besides the sequences, IMGT also annotates the start position of CDR3 in the V gene (104^th^ amino acid of the V gene in IMGT coordination). IMGT also defines the end position of CDR3 as amino acid W/F in the amino acid motif W/FGxG in the J gene. Finally, TRUST4 determines the CDR3 coordinate based on these IMGT conventions after the identification of V and J genes (9).

---

**Figure S1.**
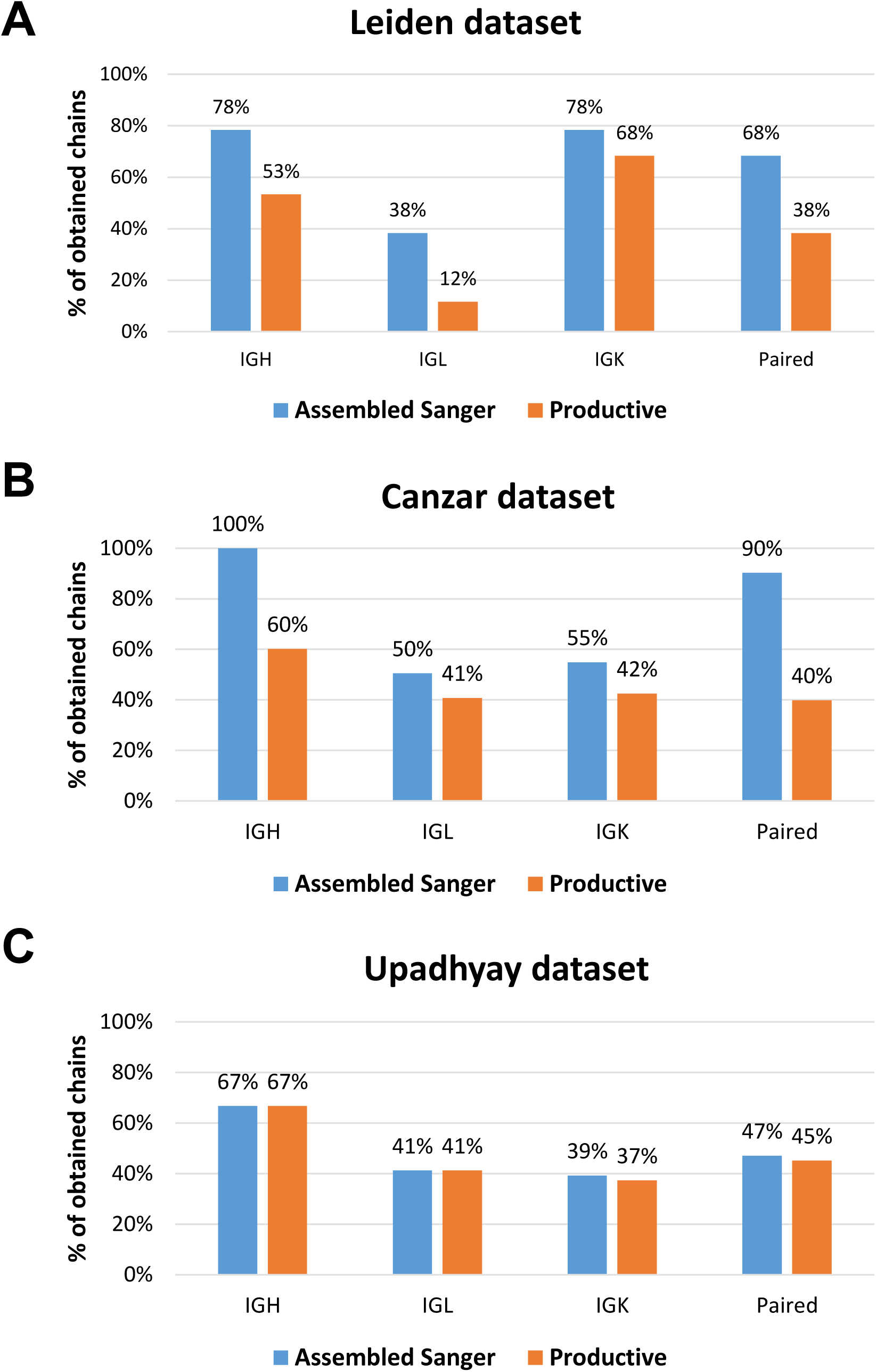
Distribution of assembled and productive heavy, light and paired (heavy and light) chains in the ground truth of each dataset. **(A)** Dataset generated in this work (Leiden) consisting of 72 antigen-specific memory B cells. **(B)** Dataset from Canzar et al. 2017 study consisting of 113 plasmablast cells. **(C)** Dataset from Upadhyay et al. 2018 study consisting of 51 plasmablast cells. *Assembled Sanger* are heavy or light chains without a stop codon, obtained after running IgBLAST on fasta files generated using Sanger sequencing. *Productive* are assembled heavy or light chains with in-frame V-J junctions.

**Figure S2.**
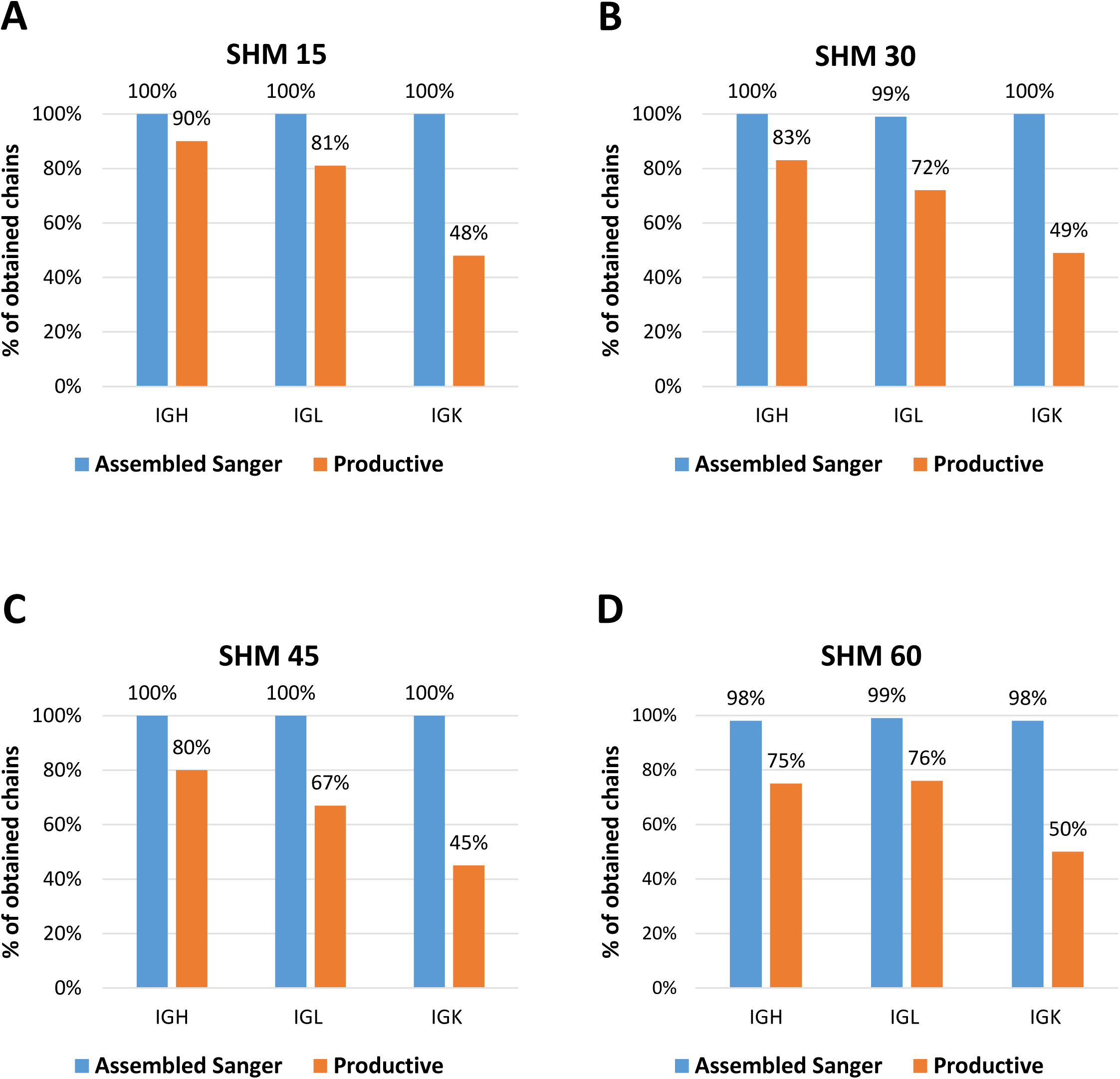
Distribution of 100 assembled and productive heavy and light chains in simulated synthetic datasets with different amounts of somatic hypermutations (SHMs). **(A)** 15 SHMs, **(B)** 30 SHMs, **(C)** 45 SHMs, **(D)** 60 SHMs. *Assembled Sanger* are heavy or light chains without a stop codon, obtained after running IgBLAST on fasta files generated by immuneSIM. *Productive* are assembled heavy or light chains with in-frame V-J junctions. The sequences of these four simulations were used as the ground truth to evaluate each tool.

**Figure S3.**
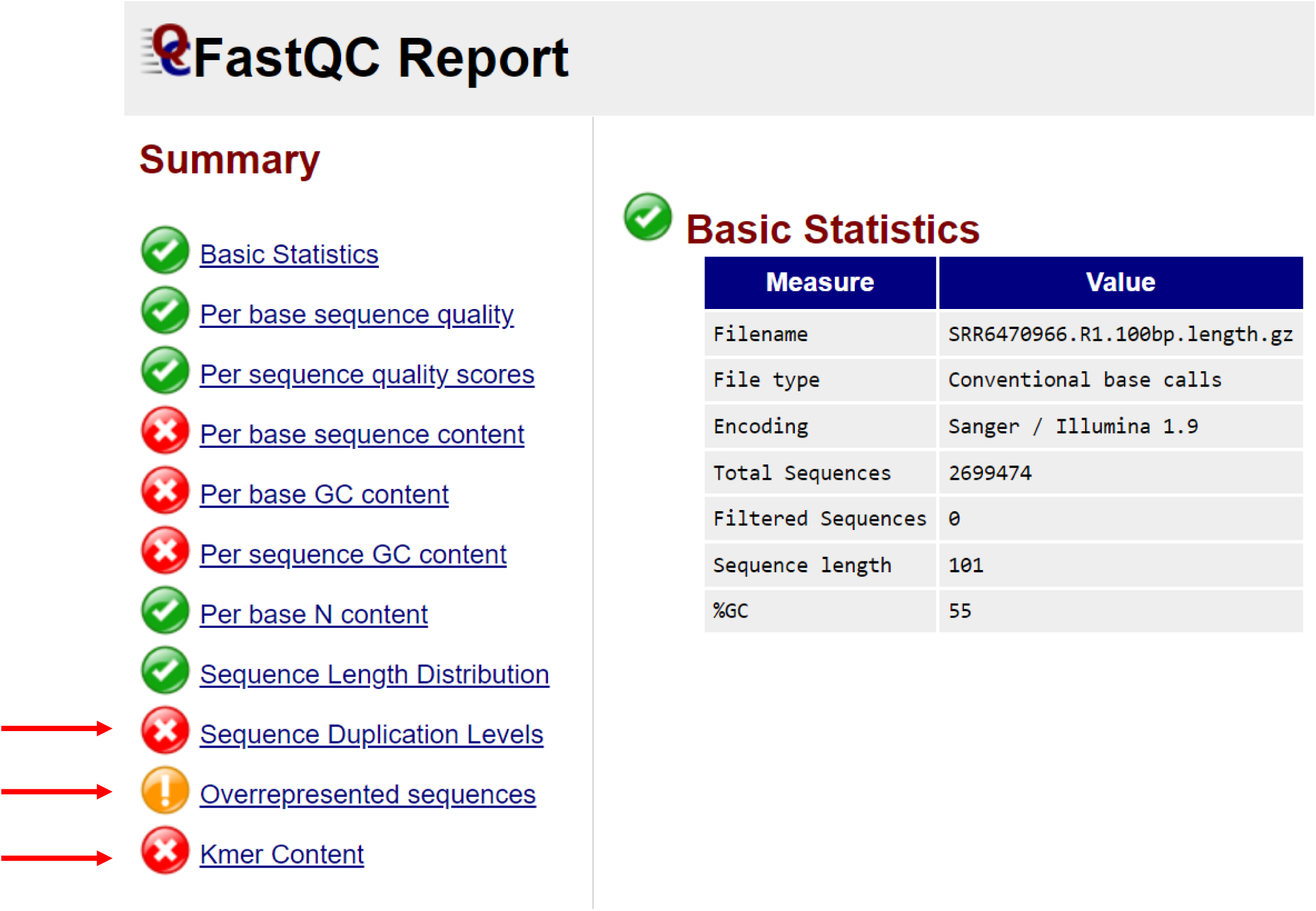
Presence of *k*-mers and overrepresented sequences in Upadhyay dataset. Quality control results of raw reads of a representative sample SRR6470966 generated using fastQC. The complete quality control results of all samples can be found in the Supplementary Data 1: fastqc_report.html

**Figure S4.**
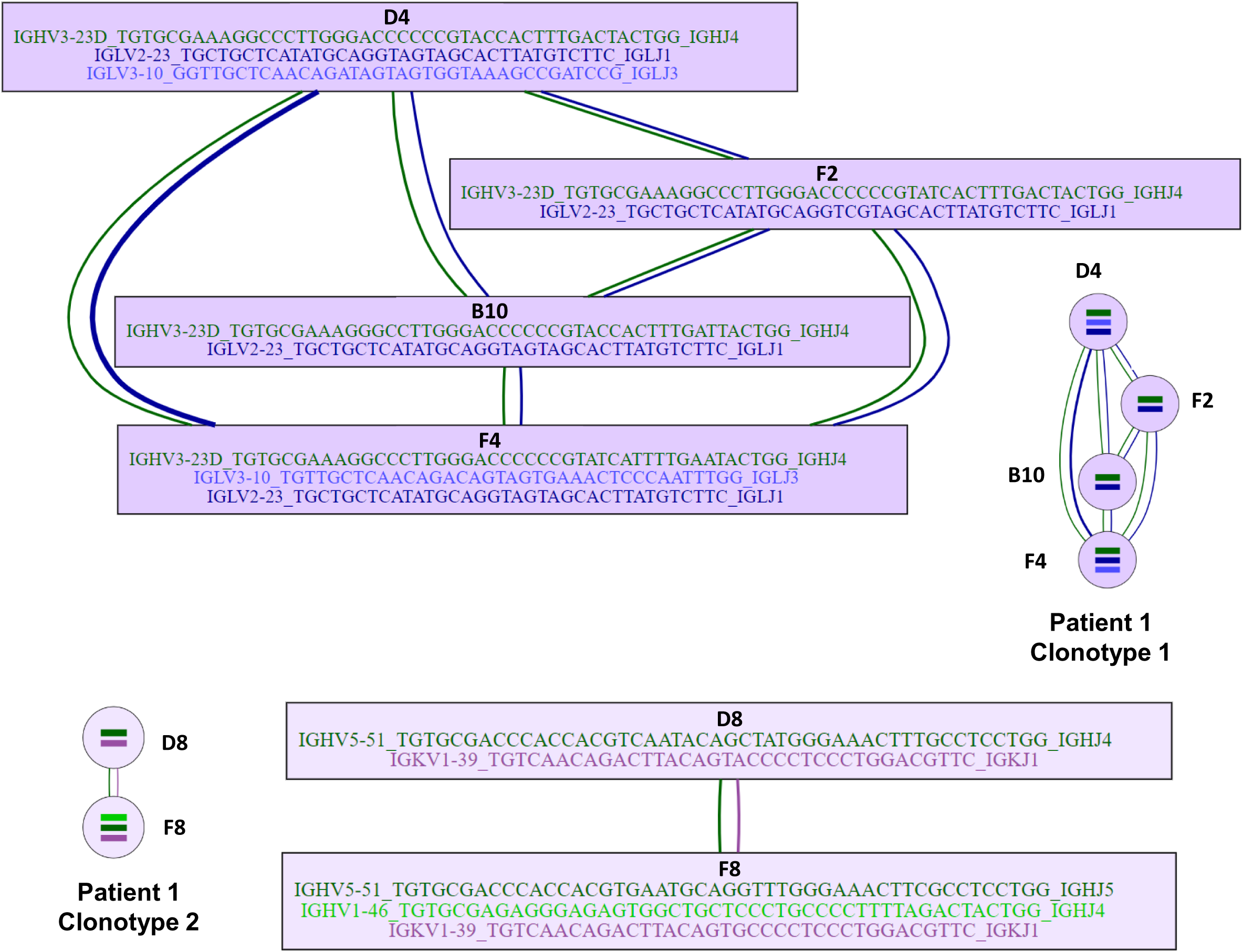
Clonotype network reconstruction. Two patient-specific BCR clonotypes, consisting of four and two cells, that were reconstructed using tetanus toxoid-specific (TT+) single cells by BRACER.

**Figure S5.**
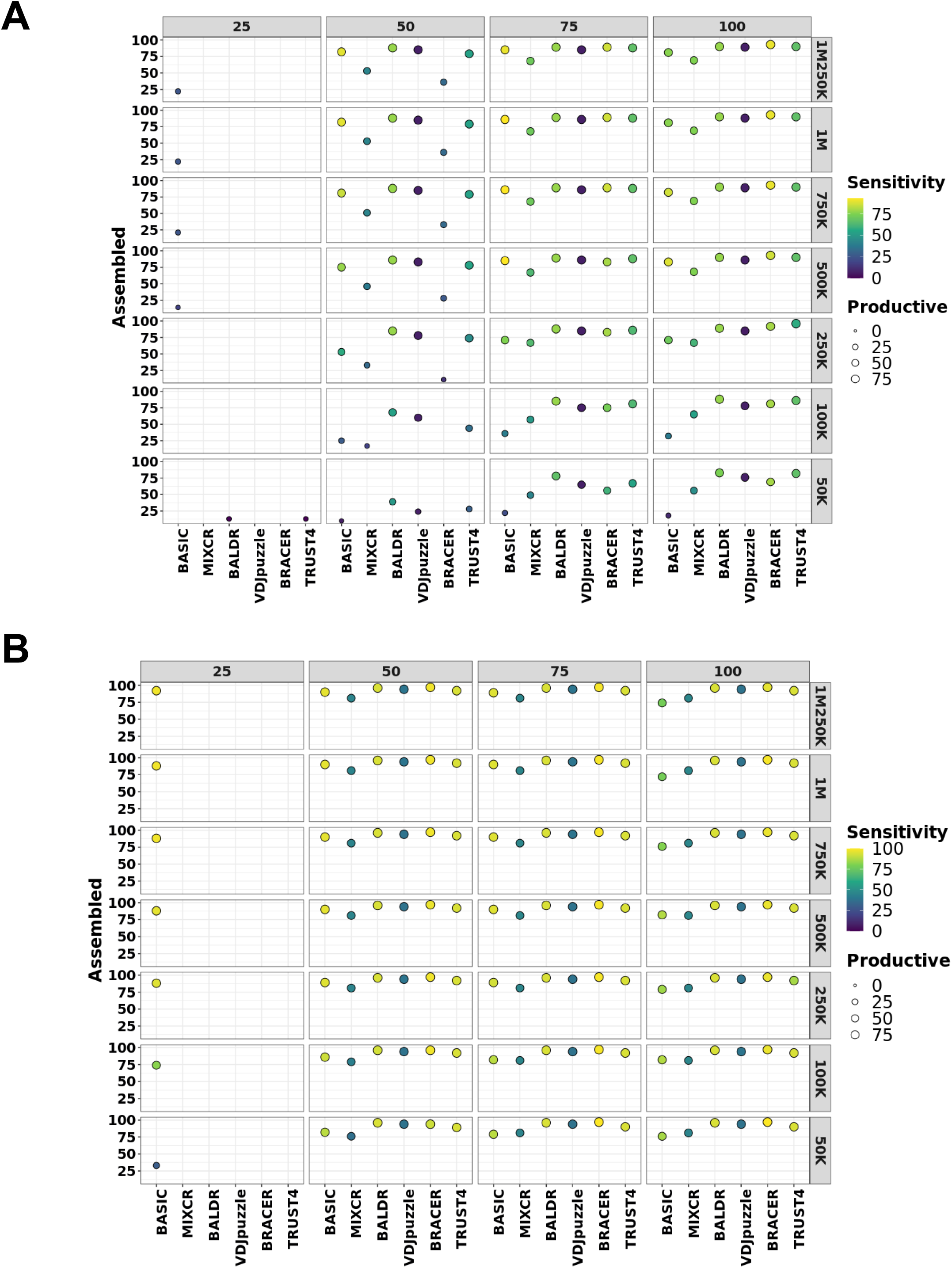
Performance of each method on Leiden dataset. Effect of different read length and coverage on the assembly, productivity and sensitivity of **(A)** Heavy chain and **(B)** Light chain. *Assembled* are heavy or light chains without stop codons. *Productive* are assembled heavy or light chains with in-frame V-J junctions. Left y-axis depicts % of assembled chains over the total number of single cells in the dataset. Right y-axis corresponds to the coverage. The size of the circles is proportional to the % of productive chains. Higher intensity of the yellow color of the circles corresponds to the higher sensitivity.

**Figure S6.**
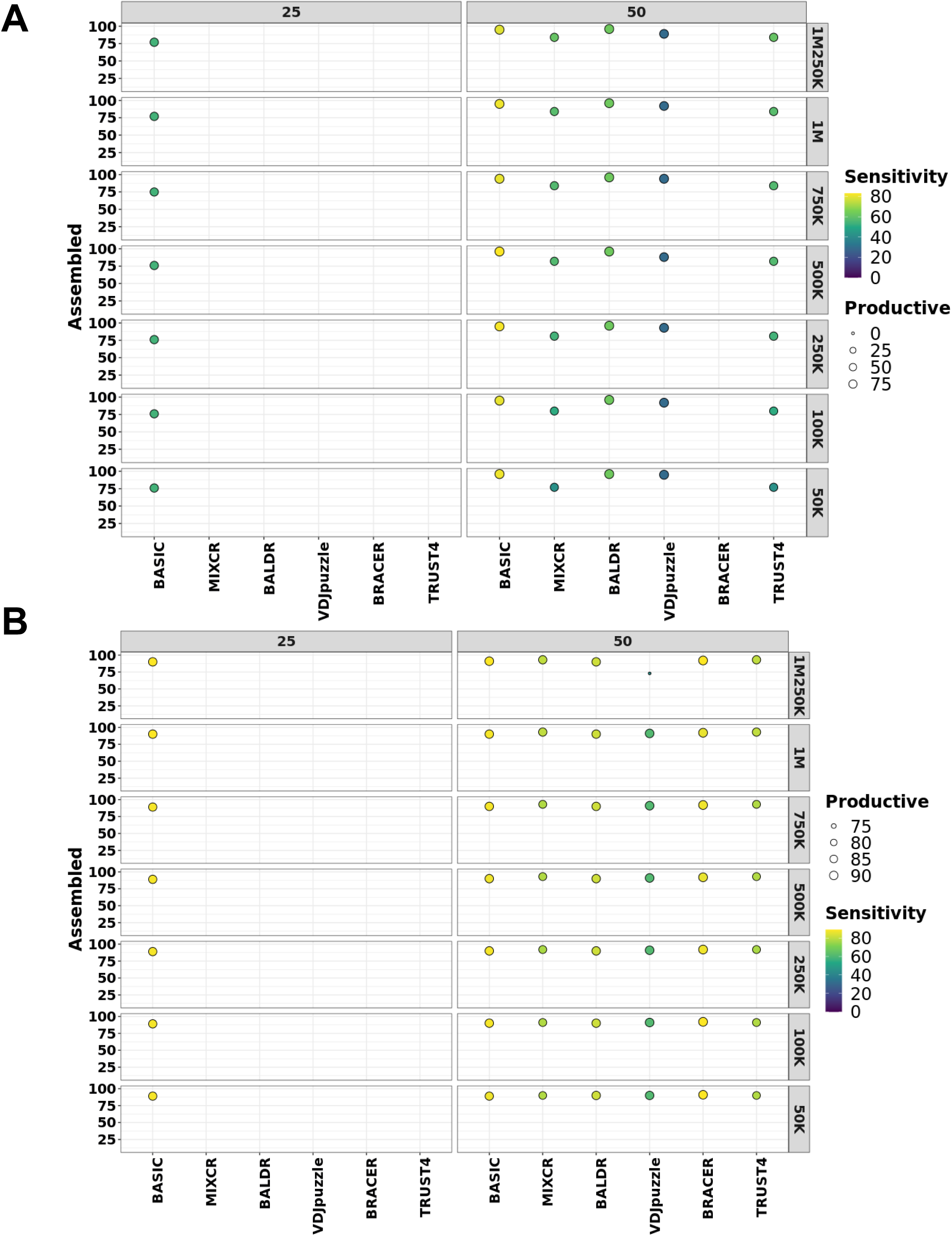
Performance of each method on Canzar dataset. Effect of different read length and coverage on the assembly, productivity and sensitivity of **(A)** Heavy chain and **(B)** Light chain. *Assembled* are heavy or light chains without stop codons. *Productive* are assembled heavy or light chains with in-frame V-J junctions. Left y-axis depicts % of assembled chains over the total number of single cells in the dataset. Right y-axis corresponds to the coverage. The size of the circles is proportional to the % of productive chains. Higher intensity of the yellow color of the circles corresponds to the higher sensitivity.

**Figure S7.**
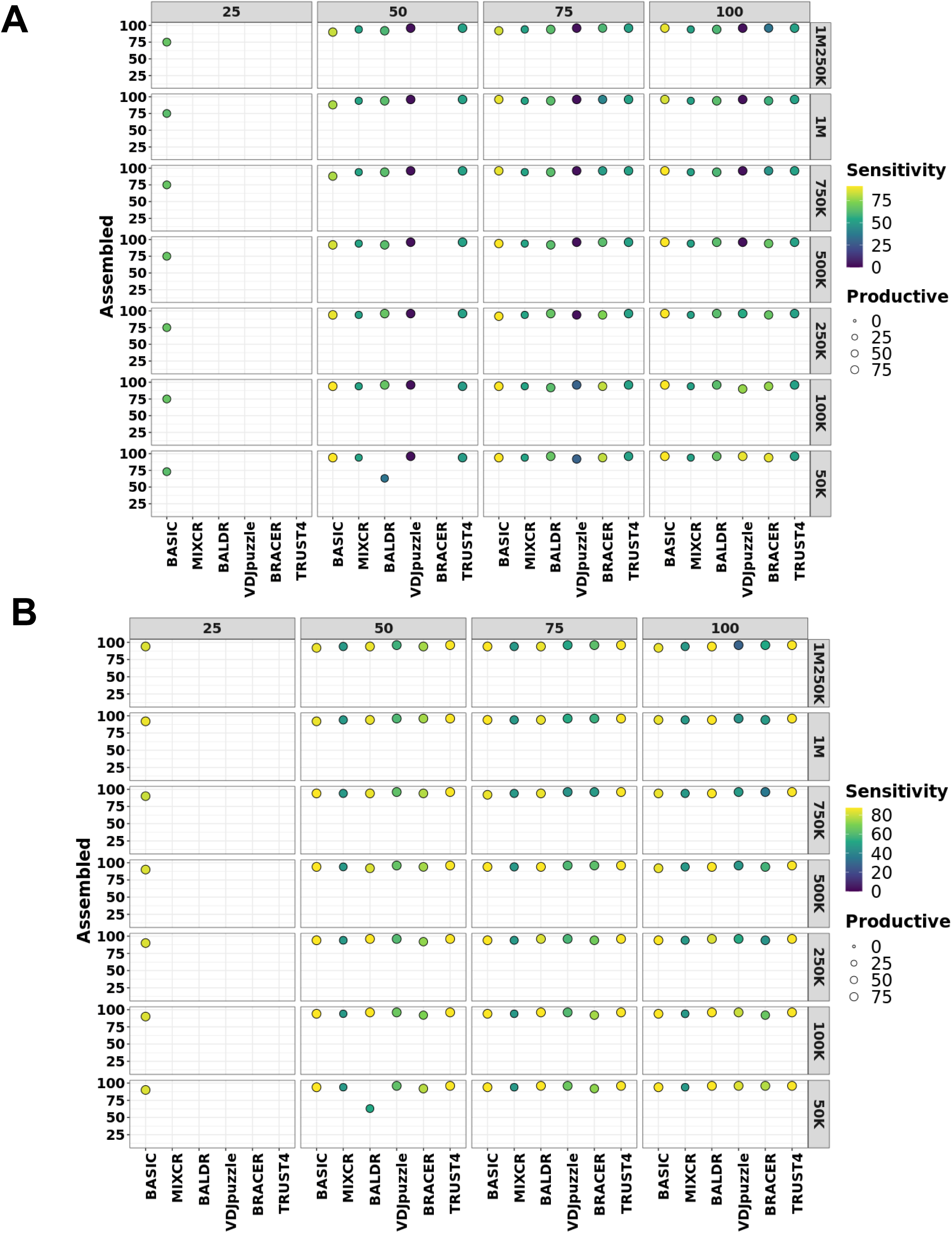
Performance of each method on Upadhyay dataset. Effect of different read length and coverage on the assembly, productivity and sensitivity of **(A)** Heavy chain and **(B)** Light chain. *Assembled* are heavy or light chains without stop codons. *Productive* are assembled heavy or light chains with in-frame V-J junctions. Left y-axis depicts % of assembled chains over the total number of single cells in the dataset. Right y-axis corresponds to coverage. The size of the circles is proportional to the % of productive chains. Higher intensity of the yellow color of the circles corresponds to the higher sensitivity.

**Figure S8.**
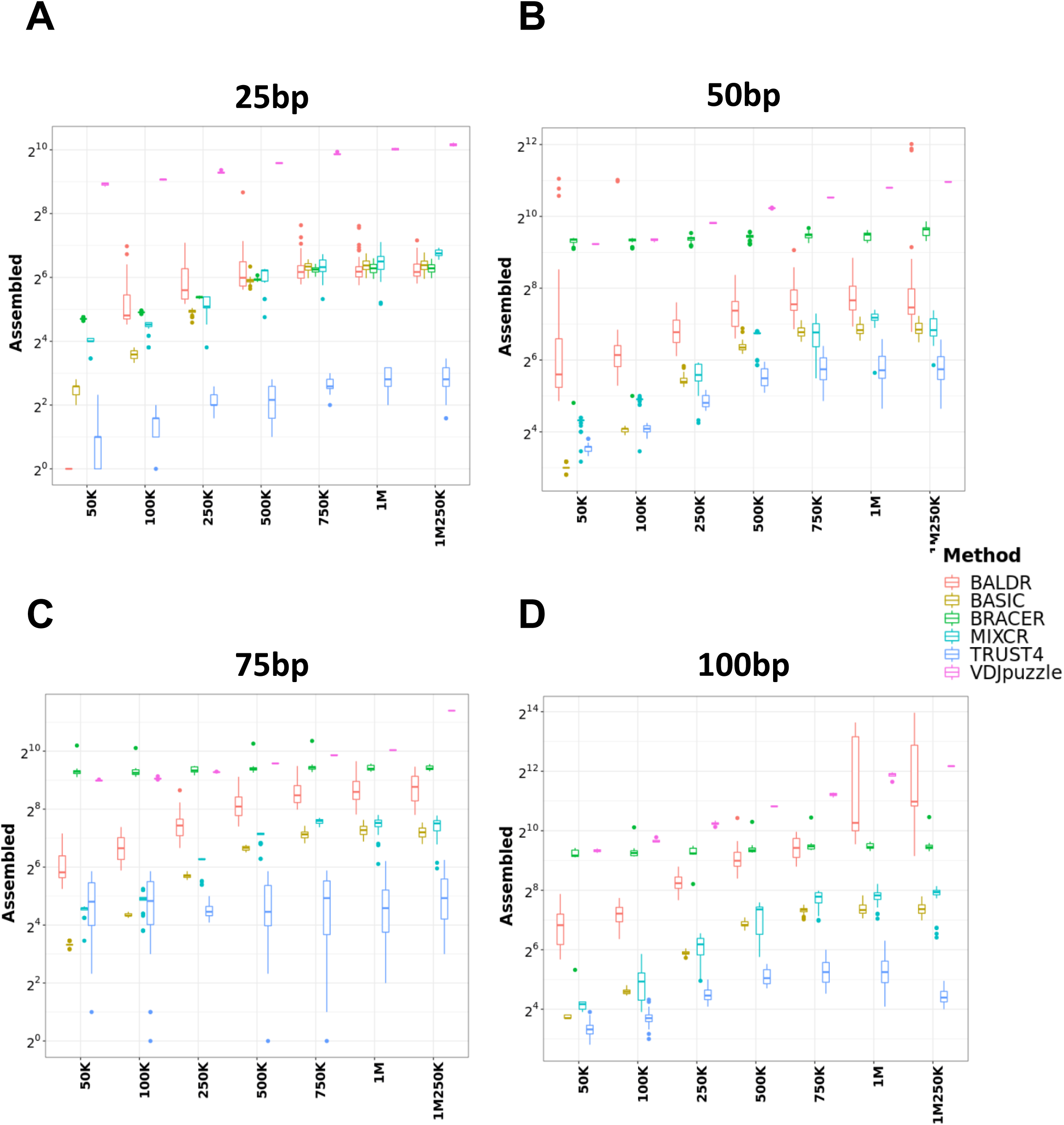
Execution time of each method using Leiden libraries with different read length and coverage: **(A)** 25 bp, **(B)** 50 bp, **(C)** 75 bp, **(D)** 100 bp. Boxplots represent interquartile ranges of the execution time for all the samples in the respective libraries. x-axis indicates the different coverage levels (from 50,000 up to 1,250,000), y-axis depicts the time in seconds, needed to process one sample.

